# Characterizing Adult cochlear supporting cell transcriptional diversity using single-cell RNA-Seq: Validation in the adult mouse and translational implications for the adult human cochlea

**DOI:** 10.1101/742270

**Authors:** Michael Hoa, Rafal Olszewski, Xiaoyi Li, Ian Taukulis, Alvin DeTorres, Ivan A. Lopez, Fred H. Linthicum, Akira Ishiyama, Daniel Martin Izquierdo, Robert J. Morell, Matthew W. Kelley

## Abstract

Hearing loss is a problem that impacts a significant proportion of the adult population. Cochlear hair cell loss due to loud noise, chemotherapy and aging is the major underlying cause. A significant proportion of these individuals are dissatisfied with available treatment options which include hearing aids and cochlear implants. An alternative approach to restore hearing would be to regenerate hair cells. Such therapy would require recapitulation of the complex architecture of the organ of Corti, necessitating regeneration of both mature hair cells and supporting cells. Transcriptional profiles of the mature cell types in the cochlea are necessary to can provide a metric for eventual regeneration therapies. To assist in this effort, we sought to provide the first single-cell characterization of the adult cochlear supporting cell transcriptome. We performed single-cell RNA-Seq on FACS-purified adult cochlear supporting cells from the *Lfng^EGFP^* adult mouse, in which supporting cells express GFP. We demonstrate that adult cochlear supporting cells are transcriptionally distinct from their perinatal counterparts. We establish cell type-specific adult cochlear supporting cell transcriptome profiles, and we validate these expression profiles through a combination of both fluorescent immunohistochemistry and *in situ* hybridization co-localization and qPCR of adult cochlear supporting cells. Furthermore, we demonstrate the relevance of these profiles to the adult human cochlea through immunofluorescent human temporal bone histopathology. Finally, we demonstrate cell cycle regulator expression in adult supporting cells and perform pathway analyses to identify potential mechanisms for facilitating mitotic regeneration (cell proliferation, differentiation, and eventually regeneration) in the adult mammalian cochlea. Our findings demonstrate the importance of characterizing mature as opposed to perinatal supporting cells.

## Introduction

Hearing loss impacts approximately 432 million adults worldwide (WHO Media centre 2018). For most of these individuals, the underlying cause of the auditory dysfunction is a loss of mechanosensory hair cells (HCs) in the cochlea. Existing data for humans, and all other mammals, strongly suggests that loss of cochlear hair cells in adults is permanent (Doetzlhofer et al. 2006; White et al. 2006). In contrast, some non-mammalian vertebrates, such as avian species, are able to robustly and repeatedly regenerate hair cells in response to injury. In these species, supporting cells (SCs), which surround the hair cells, act as precursors which give rise to new hair cells through both proliferative and non-proliferative mechanisms (Corwin and Cotanche 1988; Ryals and Rubel 1988; White et al. 2006; Liu et al. 2012; Warchol 2011; Atkinson et al. 2015).

Studies utilizing both viral gene delivery and genetically-inducible mouse models have demonstrated some success in converting supporting cells into hair cells in perinatal, but not adult, mouse models (Liu et al. 2012, 2014; Kelly et al. 2012; Kuo et al. 2015; Praetorius et al. 2010; Staecker et al. 2014). An exception to this is the recent work from Walters and colleagues demonstrating the formation of new hair cells from adult cochlear supporting cells using a combination of forced expression of *Atoh1*, a hair cell master regulator gene, and deletion of *Cdkn1b*, a gene that controls cell cycle exit in supporting cells and hair cell precursors (Walters et al. 2017). But even in this case the hair cells that were generated remained immature and, in some cases, showed signs of apoptotic cell death (Walters et al. 2017). These results suggest that our understanding of the genetic pathways that must be modulated to achieve a biological restoration of hearing remains limited.

While considerable effort has been devoted to understanding the genetic pathways that modulate hair cell formation, it is equally important to determine how cells are specified as supporting cells. As discussed, regenerated hair cells will, most likely, be derived from supporting cells, which will require de- and then re-differentiation potentially similar to the process observed in avian species (Stone and Cotanche 2007). The ability to accomplish supporting cell-mediated hair cell regeneration in the adult organ of Corti is limited by barriers that remain to be characterized. A first step towards overcoming these barriers would be to generate expression profiles for specific supporting cell types within the adult cochlea. Therefore, we sought to characterize the transcriptomes of adult cochlear supporting cells using single cell RNA-sequencing. The results demonstrate that adult cochlear supporting cells are transcriptionally distinct from their perinatal counterparts. We identify adult cochlear supporting cell-specific expression profiles and extensively validate these expression profiles through a combination of fluorescent immunohistochemistry and *in situ* hybridization co-localization in adult cochlear cross-sections and qPCR from isolated adult cochlear supporting cells. To examine the relevance of these pathways for potential clinical applications, we demonstrate expression of several novel, cell-type specific markers using immunofluorescence on human temporal bones. Finally, we perform cell cycle pathway analyses on FACS-purified single adult supporting cell transcriptomes to explore potential mechanisms to overcome adult supporting cell quiescence.

## METHODS

## EXPERIMENTAL MODEL AND SUBJECT DETAILS

## METHOD DETAILS

### Mice

CD-1 mice were obtained from Charles River Laboratories. CBA/J mice were obtained from Jackson Laboratories. The Tg(Lfng-EGFP)HM340Gsat BAC transgenic mouse line (Lfng*^EGFP^*) was generated by the GENSAT Project (Gong et al. 2003) and was kindly provided by A. Doetzlhofer (Johns Hopkins University). P60 and P120 mice of either sex were used for all experiments. Four mice were utilized for each experiment. All experiments were conducted in accordance with NIH animal use protocol 1379-15.

### Adult Cochlea Preparation

Cochleae from P60 and P120 *Lfng^EGFP^* mice were used for FACS purification of GFP+ supporting cells (SCs) before single cell capture. Apical to basal distribution of GFP+ supporting cells in the adult cochlea were determined by whole mount and orthogonal immunohistochemistry (Supplemental Figure S1, S2). Briefly, the inner ear was dissected from adult mice and the vestibular portions of the inner ear, including utricle, saccule and semicircular canal ampulla, were removed. Cochleae were then collected in a 1.5-ml tube (n = 4-5 cochleae per integrated fluidics chip (IFC) capture) and incubated in 0.05% crude trypsin (Worthington) in CMF-PBS (Life Technologies) at 37°C for 8 min. Excess trypsin solution was removed and four volumes of 5% FBS (Thermo Fisher Scientific) in DMEM/F12 (Thermo Fisher Scientific) was added to stop the digestion. The tissue was then triturated for 2 min and passed through a 40-µm strainer (pluriSelect Life Science) to eliminate residual aggregates and bone fragments. The resulting single-cell suspension was then stained with propidium iodide (Life Technologies) to allow for exclusion of dead cells and debris from the samples.

### Flow Cytometry and Sample Collection

Single cells were sorted on a FACSAria flow cytometer (BD Biosciences) with a compensated FITC setting and 488 nm excitation, using a 100-µm nozzle. In adult tissue, GFP+ SCs are the brightest population and typically comprised 2-7% of viable cells. The purpose of FACS purification was to enrich for a population of GFP+ SCs and gating was set based on wild type isotype controls to exclude the majority of GFP– cochlear cells. Scatter discrimination was used to eliminate doublets and samples were negatively selected with propidium iodide to remove dead cells. Cells were collected in 20% FBS in DMEM/F12 and stored on ice. After sorting, cells were spun at 200g for 10 min and then resuspended in 20% FBS. More detailed descriptions of the isolation methods have been previously published (Burns et al. 2015).

### Adult Cochlear Supporting Cell RNA-Seq on Fluidigm C1 Platform

Methodology for single cell RNA-Seq on the Fluidigm C1 Platform was previously described (Burns et al. 2015). Briefly, cell capture, lysis, SMARTer-based RT and PCR amplification of cDNA was performed as outlined in the Fluidigm protocol (PN 100-5950 B1). After obtaining a single-cell suspension, 10 µl of cells at a final concentration of 2.5 x 10^5^ – 7 x 10^5^ cells per µl were loaded onto a medium-sized (10-17 µm) IFC. Cell concentration was estimated at a 1:10 dilution using an automated cell counter (Luna). The IFC was placed in the C1 system, where cells were automatically washed and captured. After capture, the chip was removed from the C1 and a 30-µm stack of widefield fluorescence and brightfield images was recorded at each capture site using an x 10/0.4 numerical aperture objective on an inverted Zeiss Axio Observer.Z1 microscope equipped with a motorized stage (example image in Supplementary Figure S3). Automated imaging was performed using a custom script written in the Zeiss Zen Blue software as described previously (Burns et al. 2015). Average imaging time for all 96 capture sites was 35 min. A summary of each C1 capture can be found in Supplementary Table 1. After the imaging period, the IFC was returned to the C1 where lysis, RT and PCR were performed automatically within individual reach chambers for each cell. For RNA-Seq, mixes were prepared from the SMARTer Ultra Low RNA kit (Clontech) according to the volumes indicated in the Fluidigm protocol. For qPCR, mixes were prepared from the Single Cell-to-CT qRT-PCR kit (Ambion). The thermal cycler within the C1 performs 21 or 18 rounds of PCR amplification to obtain enough material for RNA-Seq or qPCR, respectively. cDNA was manually collected from the output channel of each capture site and stored in a 96-well plate at –20°C until library preparation. The average time from dissection to cell lysis was approximately 4 hours (h) for FACS-purified cochlear SCs.

### Fluidigm C1-generated single-cell RNA-Seq library sequencing, alignment, and estimation of gene expression

Each collection of 96 pooled single cell libraries was sequenced on a single flow cell lane of an Illumina HiSeq 1500 to an average depth of 1.8M reads using 90x90 paired-end reads. A total of 279 single cell libraries were generated from FACS-purified P60 and P120 Lfng^EGFP^-positive adult cochlear supporting cells. The quality and read depth requirements of the library preparation and sequencing protocols used here have been described in detail elsewhere, and our own analyses were all in agreement with these published reports (Burns et al. 2015).

Reads were de-multiplexed and then aligned to a Bowtie index genome based on the NCBI-annotated mouse transcriptome (comprising 48,714 genes in GRCm38.vM11 genome and corresponding GTF) using STARv2.5. The sequences and identifiers for EGFP were appended to the genome FASTA and the GTF prior to creating the index used for alignment. For each cell (library), absolute counts were calculated using STARv2.5 and using the TranscriptomeSAM parameter, relative transcript abundances were estimated from the aligned reads using RSEM v1.2.19 (default parameters) (Li and Dewey 2011; Li et al. 2010). RSEM estimates transcript abundance in units of transcript per million (TPM). The abundances reported here are at the gene level, which RSEM calculates by summing the estimated transcript abundances for each gene. Alignment, absolute counts, and abundance estimation were carried out on the NIH/Helix Biowulf cluster. Quality metrics were calculated using the CollectRNASeqMetrics in Picard. Subsequent analysis has been performed with the dataset in absolute count format. Data in TPM format is provided for comparison to previously deposited FACS-purified P1 cochlear supporting cells (Burns et al. 2015)(GEO Accession ID: GSE71982).

### Outlier identification

Cells that appeared unhealthy, as noted by lack of GFP or fragmented cellular appearance in the recorded capture site images, were excluded from library preparation. To further identify potentially unhealthy cells with abnormally low expression levels, we passed the cells through the outlier identification function provided in SINGuLAR Analysis Toolset 3.5.2, Fluidigm’s R package for single-cell expression analysis. Outlier identification in SINGuLAR proceeds by trimming low-expressing transcripts until 95% of the transcripts that remain are above 1 nTPM in half of the cells. A distribution of combined transcript expression is created from these cells, and outliers are considered as cells whose median expression across the identified gene list is below the 15^th^ percentile of the distribution. Initial validation of clustering and close examination of whole mount adult Lfng-EGFP mice cochleae revealed an additional population of Lfng^EGFP^-positive cells which constituted endothelial cells of capillaries in the spiral ganglion region. This along with the previously published outlier analysis methodology resulted in a final dataset of 211 adult cochlear supporting cells after the exclusion of outliers. Using these routines, 68 out of 279 adult cochlear supporting cells were excluded from the analysis (Supplementary Table S1).

### PCA and *t*-SNE Analysis

#### Selection of genes for clustering analysis

Identification of the highly variable genes was performed in Seurat utilizing the MeanVarPlot function and the default settings with the aim to identify the top ∼ 2000 variable genes (Satija et al. 2015). Briefly, to control for the relationship between variability and average expression, average expression and dispersion is calculated for each gene, placing the genes into bins, and then a z-score for dispersion within the bins was calculated. These genes are utilized in the downstream analyses for clustering.

#### Clustering of single cells

Clustering analyses of single-cell data was performed with Seurat using a graph-based clustering approach (Satija et al. 2015). Briefly, the Jackstraw function using the default settings was used to calculate significant principal components (p < 0.0001) and these principal components were utilized to calculate k-nearest neighbors (KNN) graph based on the Euclidean distance in PCA space. The edge weights are refined between any two cells based on the shared overlap in their local neighborhoods (Jaccard distance). Cells are then clustered according to a smart local moving algorithm (SLM), which iteratively clusters cell groupings together with the goal to optimize the standard modularity function (Blondel et al. 2008; Waltman and van Eck 2013)(http://satijalab.org/seurat/pbmc-tutorial.html). Resolution in the FindClusters function was set to 0.8. High modularity networks have dense connections between the nodes within a given module but sparse connections between nodes in different modules. Clusters were then visualized using a t-distributed stochastic neighbor embedding (t-SNE) plot.

#### Differential gene expression analysis

Differential expression analysis was performed in Seurat utilizing the FindAllMarkers function with the default settings except that the “min.pct” and “thresh.use” parameters were utilized to identify broadly expressed (min.pct = 0.8, thresh.use = 0.01) and subpopulation-specific (min.pct = 0.5, thresh.use = 0.25) expression profiles. The parameter “min.pct” sets a minimum fraction of cells that the gene must be detected in all clusters. The parameter “thresh.use” limits testing to genes which show, on average, at least X-fold difference (log-scale) between groups of cells. The default test for differential gene expression is “bimod”, a likelihood-ratio test (McDavid et al. 2013). Differentially expressed genes were then displayed on violin plots based on unbiased clustering described above. Default parameters for significance included a minimum of 25% of cells expressing a given gene and a fold change of at least 1.7.

#### Validation of qPCR primer set assays

A total of 96 DELTAgene qPCR gene expression assays, consisting of forward and reverse qPCR primers, were purchased from Fluidigm. 93 of the 96 assays were used for single-cell qPCR analysis in this study. DELTAgene assays were validated against adult mouse cochlea cDNA. To determine primer efficiencies for all 96 primer sets, we made 8 threefold dilutions of the preamplification product and tested them in triplicate on a 96.96 Dynamic Array IFC using a Fluidigm BioMark HD microfluidics-based qPCR system. Primer efficiency was calculated with the formula: E = 10 (-1/slope), where slope represents the linear slope of a linear regression fit to the average standard curve. Primer sets with efficiencies ± 0.2 from the ideal efficiency of 2, or primer sets where multiple peaks were detected on a melt curve were excluded from further analysis.

### Validation of adult cochlear supporting cell gene expression with single cell qPCR and digital droplet PCR (ddPCR)

#### Single-cell qPCR

For single-cell qPCR, we captured single cells from P60 cochlea using medium-sized microfluidic chips (C1 Single-Cell Auto Prep IFC for PreAmp) as outlined in the Fluidigm protocol (PN 100-6117 G1). Mixes for lysis, RT, and specific target amplification were prepared from the Single Cell-to-Ct qRT-PCR kit (Ambion, Austin, TX) and pre-designed Delta Gene assays (Fluidigm, South San Francisco, CA). In addition to known markers, putative adult cochlear supporting cell markers were selected for analysis in an arbitrary manner.

After 18 cycles of preamplification, expression levels using single cell qPCR were performed on the Fluidigm Biomark HD system as previously described (Honda et al. 2017). cDNA from single cells was selected for qPCR in the same way as it was selected for RNA-Seq. A total of 170 single cells from four C1 captures were profiled using 2 Dynamic Array IFCs. Empty wells with primers were utilized for negative controls. The threshold of cycles (C_t_) values were calculated with Fluidigm Real-time PCR analysis software with the following settings: quality threshold of 0.65; a linear (derivative) baseline correction; and auto (detectors) method. We defined gene expression levels as log_2_ expression = LOD – C_t_, in which C_t_ = 24 was set as the LOD. We used the log_2_ expression dataset for hierarchical clustering. The SINGuLAR package v3.6.1 was utilized to display and analyze single-cell qPCR data.

#### Digital droplet PCR (ddPCR)

FACS-purified adult cochlear supporting cells were collected from P60 Lfng*^EGFP^* mice as detailed previously. RNA was isolated from 30,000 GFP-positive and GFP-negative cells with each assay standardized to 2,000 cells per sample collected in triplicates to use for quantification experiment. ddPCR Droplets were generated using the QX200 AutoDG Droplet Digital. PCR was performed as described in the QX200™ ddPCR™ EvaGreen® Supermix instructions (http://www.bio-rad.com/webroot/web/pdf/lsr/literature/10028376.pdf). Droplets were read with a QX200™ Droplet Reader (BioRad) and analyzed with QuantaSoft software (BioRad).

#### Co-localization with immunofluorescence and single molecule fluorescent *in situ* hybridization (smFISH)

Cochleae were fixed in fresh 4% paraformaldehyde in PBS overnight at 4°C. After fixation, specimens were washed in PBS then permeabilized and blocked for 1 hour at room temperature in PBS with 0.2% Triton X-100 (PBS-T) with 10% fetal bovine serum. Samples were then incubated in the appropriate primary antibodies overnight at 4°C in PBS-T with 10% fetal bovine serum, followed by three rinses in PBS-T and labelling with AlexaFluor-conjugated secondary antibodies (1:250, Life Technologies) in PBS-T for 1 hour at room temperature. Where indicated, 4,6-diamidino-2-phenylindole (DAPI) (1:10,000, Life Technologies) was included with the secondary antibodies to detect nuclei. Organs were rinsed in PBS three times and mounted in SlowFade (Invitrogen). Specimens were imaged using a Zeiss LSM 710 confocal microscope.

For immunohistochemistry and *in situ* hybridization of cochlear sections, fixed adult mouse inner ears were decalcified in 150 mM EDTA for 5-7 days, transferred to 30% sucrose, and then embedded and frozen in SCEM tissue embedding medium (Section-Lab Co, Ltd.; Hiroshima, Japan). Adhesive film (Section-Lab Co, Ltd.; Hiroshima, Japan) was fastened to the cut surface of the sample in order to support the section and cut slowly with a blade to obtain 6 µm thickness sections. The adhesive film with section attached was submerged in 100% EtOH for 60 seconds, then transferred to distilled water. The adhesive film consists of a thin plastic film and an adhesive and it prevents specimen shrinkage and detachment. This methodology allows for high quality anatomic preservation of the specimen. Sections were cut to a thickness of 10 micrometers. Fluorescent immunohistochemistry was performed as described above. Sections were mounted with SCMM mounting medium (Section-Lab Co, Ltd.; Hiroshima, Japan).

Fluorescent *in situ* hybridization was performed using RNAscope® probes against candidate genes per previously published methodology and overlaid with fluorescent IHC (F. Wang et al. 2012). Briefly, tissue preparation was modified from the original published protocol (Advanced Cell Diagnostics, 320534) to avoid tissue detachment during the target retrieval step. Drying after sectioning was extended to 2 h followed by 1 h post-fixation in 4% PFA. A 30 min dry baking step at 60°C was added before protease treatment. Hybridization and detection were performed as described in RNAscope® 2.5 HD Detection Reagent – RED User Manual Part 2 (Document Number 322360-USM) without hematoxylin counterstaining. The following RNAscope® probes were used: human *TUBA1B* (Hs-TUBA1B), mouse *Myh9* (Mm-Myh9, 556881), mouse *Cdk1nb* (Mm-Cdkn1b, 499991), mouse *S100a6* (Mm-S100a6, 412981), mouse *Lcp1* (Mm-Lcp1, 487751), mouse *Notch2* (Mm-Notch2, 425161), mouse *Nlrp3* (Mm-Nlrp3, 439571), mouse *Slc2a3* (Mm-Slc2a3, 438851), mouse *Pla2g7* (Mm-Pla2g7, 453811), mouse *Spry2* (Mm-Spry2, 425061), mouse *Birc5* (Mm-Birc5, 422701), and mouse *Notch2* (Mm-Notch2, 425161). The target genes and probed regions are listed in Supplemental Table S4. Sequences of target probes, preamplifier, amplifier, and label probe are proprietary (Advanced Cell Diagnostics, Newark, CA). For fluorescent detection, the label probe was conjugated to Alexa Fluor 555. Assays were performed in parallel with positive (Ppib) and negative (dapB) controls.

For immunostaining following *in situ* hybridization, slides were washed 3 x 10 min in PBS-T and blocked for 3 h in 10% FBS before standard immunofluorescence procedure detailed previously (Burns et al. 2015). DAPI counterstaining was performed to label cell nuclei. The following primary antibodies were used: rabbit anti-Myosin VIIA (1:250; Proteus BioSciences, 25-6791), sheep anti-S100a6 (1:100; R&D, AF4584), rabbit anti-Lcp1 (1:100; Cell Signaling, 3588S), and mouse anti-Acetylated tubulin (1:250; Sigma). Candidate genes were tested on at least 3 adult mouse specimens from 3 different mice.

#### Human temporal bone fluorescent immunohistochemistry

Immunohistochemistry on human temporal bones was performed as previously described by Lopez and colleagues (Lopez et al. 2016). Detailed procedures of temporal bone collection, fixation, decalcification and celloidin embedding were described by Merchant (Merchant 2010). The methodology to mount celloidin-embedded sections, celloidin removal and antigen retrieval steps has been described in detail (O’Malley, Merchant, et al. 2009; O’Malley, Burgess, et al. 2009; S. R. Shi, Cote, and Taylor 1998; Huang 1975) and used previously by the authors (Lopez et al. 2016). Briefly, sections were removed from the archival jar and immediately floated in 80% ethanol and mounted on subbed glass slides (Superfrost Plus glass slides, Thermo Scientific). Bibulous paper soaked with 10% formalin was placed over the section. A small roller was used to flatten the sections. A 4” x 4” block of wood was placed over each slide. Sections were allowed to dry for 1 day, and the weight and the bibulous paper were removed. Celloidin removal was performed by immersing the sections in sodium ethoxide diluted in ethanol (1:3 ratio) for 30 minutes followed by sequential immersions in 100% acetone, methanol (100%, 70%, 50% for 5 minutes each), distilled water, 5% hydrogen peroxide (10 minutes) and then washed with distilled water prior to antigen retrieval. Antigen retrieval was performed as described previously (Lopez et al. 2016). Briefly, sections were heated in a microwave oven using intermittent heating methods of two 2-minute cycles with an interval of 2 minutes between the heating cycles in 1:200 diluted antigen retrieval solution in water (Vector Antigen Unmasking Solution, Vector Labs, Burlingame, CA, USA). The petri dish containing the slides was removed from the microwave oven and allowed to cool for 15 min at room temperature and washed with PBS for 10 min prior to immunohistochemistry. Quenching of auto-fluorescence prior to immunohistochemistry was performed as described to remove auto-fluorescence intrinsic to the human temporal bone sections (Lopez et al. 2016). Briefly, sections were placed in glass Petri dish containing ice cold PBS and placed in a UV chamber for 8 h. Temperature was checked continuously to avoid overheating, and cold PBS was replaced every 30 minutes. Sections were process for immunofluorescence once the auto-fluorescence signal in the tissue sections disappeared. Immunofluorescence was performed as previously described with hydrogen peroxide incubation step omitted. Sections were incubated with primary antibodies for 48 h at room temperature and secondary antibodies for 2 h at room temperature. The following primary antibodies were used: sheep anti-S100a6 (1:100; R&D, AF4584), rabbit anti-Lcp1 (1:100; Cell Signaling, 3588S), mouse anti-Acetylated tubulin (1:250; Sigma), and calbindin (1:100; Santa Cruz, sc-365360). Candidate genes (sheep anti-S100a6, rabbit anti-Lcp1) were tested on at least 3 human specimens each from different individuals.

#### Gene ontology, gene-set enrichment analysis and pathway-enrichment analysis of predicted proteins

Gene ontology analysis and gene enrichment analysis were performed using Enrichr (http://amp.pharm.mssm.edu/Enrichr/) as previously described (E. Y. Chen et al. 2013; Kuleshov et al. 2016; Pazhouhandeh et al. 2017). Enrichr is an integrated web-based application that includes updated gene-set libraries, alternative approaches to ranking enriched terms, and a variety of interactive visualization approaches to display the enrichment results. Enrichr employs three approaches to compute enrichment as previously described (Jagannathan et al. 2017). The combined score approach where enrichment was calculated from the combination of the p-value computed using the Fisher exact test and the z-score was utilized. In order to visualize molecular interaction networks, the list of putative proteins inferred from genes expressed by adult cochlear supporting cells was introduced to STRING 10.0 (http://string-db.org) and the nodes (proteins) and edges (protein-protein interactions) were extracted (Szklarczyk et al. 2015). Proteins were linked in STRING based on the default medium (0.400) minimum interaction score and on the following seven criteria: textmining, experiments, databases, co-expression, neighborhood, gene-fusion, and co-occurrence. Interaction evidence from all utilized criteria are benchmarked and calibrated against previous knowledge, using the high-level functional groupings provided by the manually curated Kyoto Encyclopedia of Genes and Genomes (KEGG) pathway maps (Szklarczyk et al. 2015). The summation of this interaction evidence is utilized to construct a minimum interaction score.

## QUANTIFICATION AND STATISTICAL ANALYSIS

### Experimental design

For mouse experiments, all groups consisted of age-matched mice and n represents the number of independent biological replicates. Both male and female mice were pooled for all experiments. All data were included in the analysis and thus no exclusion criteria were used. Experiments were not blinded. Sample size estimates were not performed beforehand. Details are provided in the detailed methods section.

### Statistical analysis

General statistical analysis for Figures (Bulk qPCR) were performed using *n* biological replicates as indicated in the detailed methods. Error bars indicated mean ± s.e.m., and statistical differences were assessed by a t test. Statistical analysis for single-cell RNA-Seq is described in the detailed methods.

## DATA AND SOFTWARE AVAILABILITY

All data generated in these studies were deposited in the Gene Expression Omnibus (GEO) database (GEO accession ID: GSE135703) and will be available upon publication. We are also in the process of uploading the data into the gene Expression Analysis Resource (gEAR), a website for visualization and comparative analysis of multi-omic data, with an emphasis on hearing research (https://umgear.org).

## Results

### Adult cochlear supporting cells are transcriptionally distinct from perinatal cochlear supporting cells

Unbiased clustering of the P1 and mature (P60 and P120) cochlear supporting cell transcriptomes reveals that FACS-purified adult cochlear supporting cells demonstrate broadly distinct differences from previously published FACS-purified P1 cochlear supporting cells (Burns et al. 2015) (Figure 1A). No differences in clustering or gene expression were observed between P60 and P120 cochlear supporting cells and for these reasons are grouped together as mature supporting cells for the remainder of the analyses (data not shown). Average gene expression between mature (P60 and P120) and P1 cochlear supporting cells suggests both similarities (gene expression approximating linear line in red on Figure 1B) as well as distinct differences (as shown by the number of genes that diverge between the two states) between perinatal and mature supporting cells. Supplemental Figure S4 demonstrates the degree of correlation in average gene expression as well as the large cell-to-cell variability in gene expression between P1 and mature cochlear supporting cells. The non-averaged gene expression data (Figure 1C, Figure S4A) highlight the distinct gene expression profile of mature cochlear supporting cells. Using the Seurat FeaturePlot function, the differential distribution of select known cochlear supporting cell gene expression (*Dstn, Tuba1b, Notch1, S100a1*) between P1 and mature cochlear supporting cells supports the notion that mature supporting cells are transcriptionally distinct (Herde, Friauf, and Rust 2010; Oesterle and Campbell 2009; Tannenbaum and Slepecky 1997; Saha and Slepecky 2000; Maass et al. 2015; Coppens et al. 2001) (Figure 1C). To confirm the results obtained from the single cell analysis, immunohistochemical analysis for the four known marker genes was performed in P1 and P60 cochlear sections and confirms similar distinct protein expression comparable to transcriptome differences (Figure 1D). DSTN protein expression is present in both P1 and P60 cochlear supporting cells but appears less prominent and more confined to supporting cells at P60. NOTCH1 protein is expressed in cochlear supporting cells at P1 but is absent at P60. TUBA1B protein (acetylated tubulin antibody) expression is predominantly in the pillar cells at P1 with expression noted in the peripheral axons of the spiral ganglion neurons that make contact with the hair cells. In contrast, supporting cells (pillar cells and Deiters cells) at P60 exhibit diffuse expression of TUBA1B protein. S100A1 protein is expressed in inner hair cells and cochlear supporting cells at P1 but is restricted to inner border, inner phalangeal and the outermost Deiters cell in P60. These differences in transcriptome and accompanying protein expression support the transcriptional distinctiveness of mature cochlear supporting cells from their perinatal counterparts.

**Figure 1.**
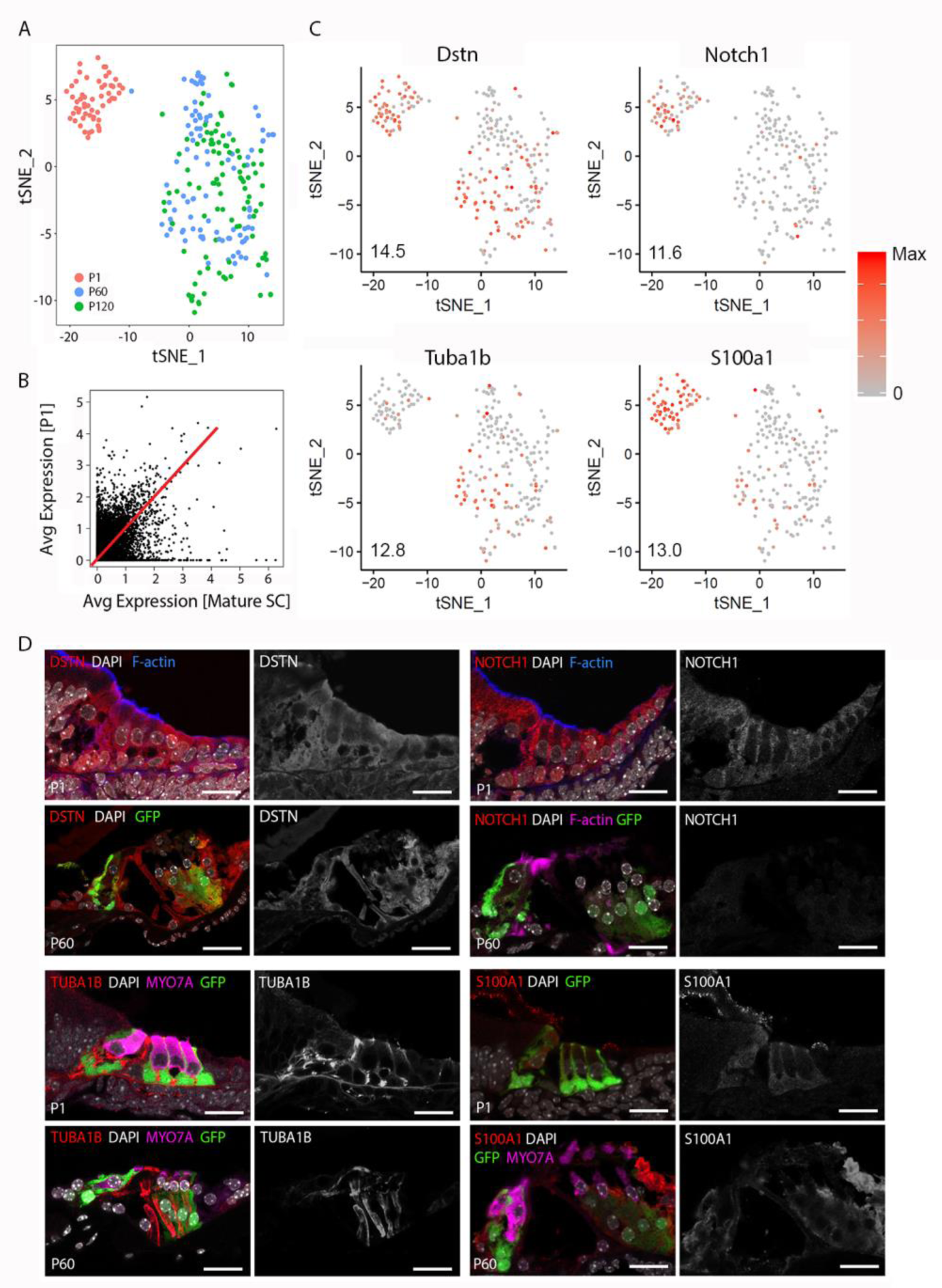
Adult cochlear supporting cells are transcriptionally distinct from perinatal cochlear supporting cells. **(A)**, Unbiased clustering of FACS-purified P1 and mature (P60, P120) cochlear supporting cell transcriptomes demonstrates clustering of single cells based on the transcriptional expression profiles for each cell. Note that P1 and mature cochlear supporting cells cluster within their respective groups but exhibit distinct clustering from each other. **(B),** Comparison of averaged gene expression between FACS-purified mature (P60, P120) and P1 cochlear supporting cells indicates both equivalent (genes expressed on or near the red line) and differential (genes located closer to either axis) expression between the two cell stages. **(C),** Feature plots of select known cochlear supporting cell genes (*Dstn, Notch1, S100a1, Tuba1b*) demonstrate distinct differences between P1 and mature cochlear supporting cells. Expression is shown in log2[nTPM] with the maximum expression value (Max) shown in the lower left corner of each plot. Expression histogram is shown with red indicating higher expression. **(D),** Representative immunohistochemistry validating transcriptional differences between P1 and mature cochlear supporting cells in organ of Corti from Lfng*^EGFP^* mice. Each 4 panel grouping demonstrates P1 and P60 immunohistochemistry with the protein of interest in the red channel (left panels) at P1 (upper left panel) and P60 (lower left panel) and the grayscale single channel images of the protein of interest (right panels) at P1 (upper right panel) and P60 (lower right panel). Known protein expression (DSTN, NOTCH1, S100A1, TUBA1B) is demonstrated (Upper left 4 panels and proceeding clockwise). Staining for F-actin or Myo7a identifies hair cell stereocilia or hair cells, respectively. Scale bar, 20 μm.

### Single-cell RNA-Seq identifies adult supporting cell gene expression

As a first step in examining this data set, expression of a subset of candidate genes was examined. A representative cross-sectional schematic of the adult *Lfng^EGFP^* mouse organ of Corti is shown in Figure 2A with GFP-expressing cells colored in green with the lighter green in pillar cells denoting less prominent GFP expression versus other supporting cell types. Initial differential expression analysis of cell clusters and close examination of whole mount adult *Lfng^EGFP^* cochleae revealed an additional population of EGFP^+^ cells that constituted endothelial cells of capillaries in the spiral ganglion region. As these cells do not constitute supporting cells, data from these cells were removed based on expression of known endothelial transcripts, and the resulting data set was subsequently filtered based on outlier analysis. The final dataset included transcriptomic values from 211 adult cochlear supporting cells. Clustering analysis identified two distinct clusters as illustrated in the heat map and tSNE plot (Figure 2B). No differences in clustering or gene expression were observed between P60 and P120 cochlear supporting cells (data not shown) and so data from the two time points were combined.

**Figure 2.**
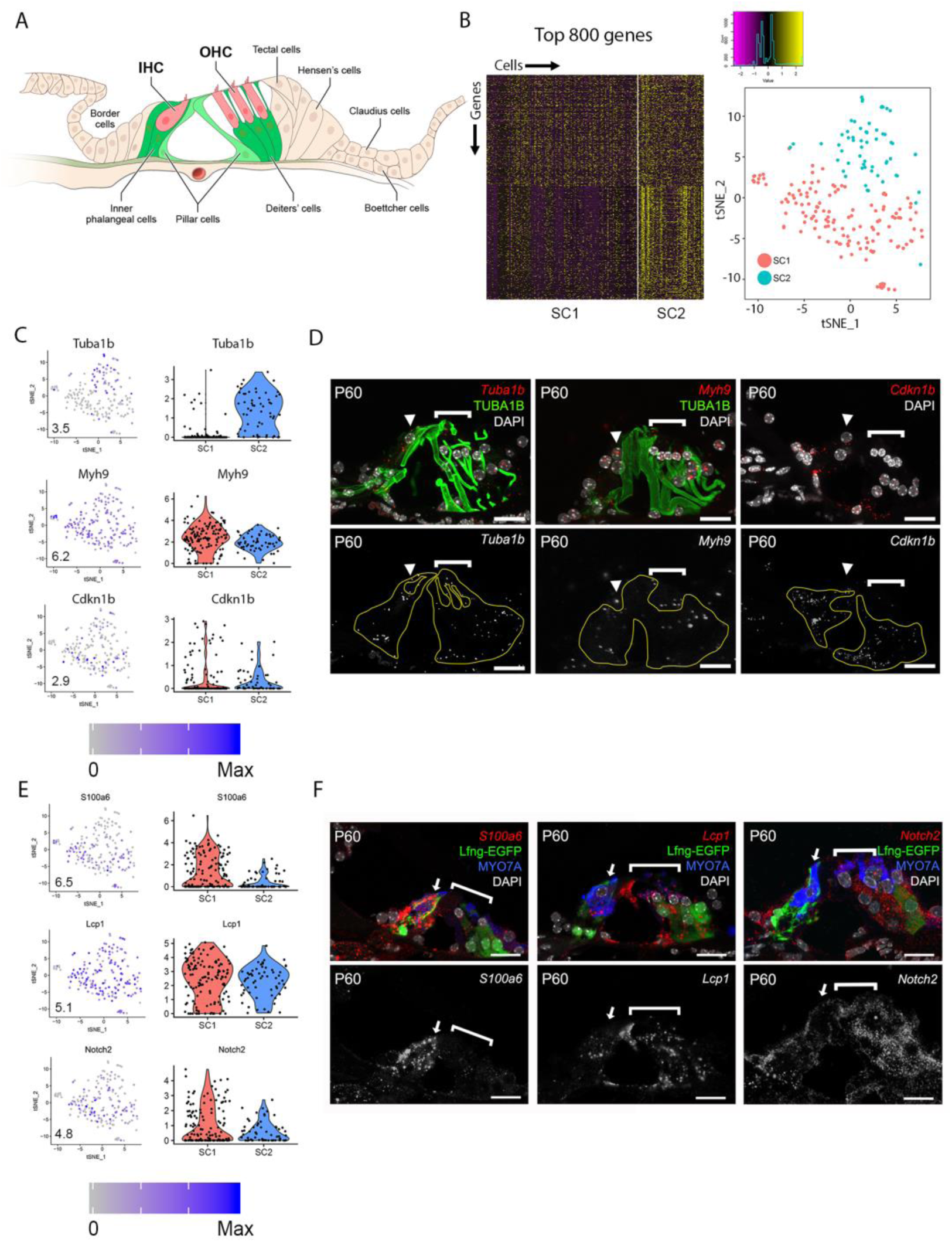
Single-cell RNA-Seq identifies adult supporting cell gene expression. **(A)**, Schematic of adult *Lfng^EGFP^*organ of Corti. GFP-expressing cells include inner phalangeal cells, Deiters’ cells, and, to a lesser extent, pillar cells. IHC = inner hair cell, OHC = outer hair cell. Mice that are homozygous and heterozygous for the transgene display the same phenotype. **(B),** Heatmap (Left) with 211 adult cochlear supporting cells (along horizontal axis) and the top 800 differentially expressed genes between adult cochlear supporting cell clusters (along vertical axis). Approximately 400 genes are localized to the upper half of the heat map while approximately 400 additional genes are localized to the lower half of the plot. tSNE plot (Right) shows two clusters of adult cochlear supporting cells (SC1 in red, SC2 in blue). **(C),** Expression of genes that are known to be expressed by adult cochlear supporting cells (*Myh9, Cdkn1b*). Left, feature plots show the expression level of each gene in each cell. Expression is shown in log2[nTPM] with the maximum expression value (Max) shown in the lower left corner of each plot. Expression histogram is shown with blue indicating higher expression. Compare with tSNE plot in Figure 2B. Right, violin plots for the same genes illustrate gene expression levels in log-scale across the two identified clusters of adult cochlear supporting cells. **(D),** smFISH localization of RNA expression for *Myh9,* and *Cdkn1b* in cross-sections of adult organ of Corti demonstrates localization in all supporting cells (border, inner phalangeal, pillar and Deiters cells). Upper left image demonstrates the *Myh9* probe in red along with antibody labeling for acetylated tubulin (TUBA1B) and DAPI-labeling of nuclei. Lower left image shows the *Cdkn1b* probe in red and DAPI-labeling of nuclei. Upper and lower panels to the right illustrates just the smFISH channel. **(E),** Expression of novel adult cochlear supporting cell genes (*Lcp1, Notch2*, *Nlrp3*, *Slc2a3*) presented as in Figure 2C. **(F),** smFISH localization of RNA expression for *Lcp1, Notch2*, *Nlrp3*, and *Slc2a3* in adult cross-sections of the organ of Corti. Images to the right demonstrate the RNA probe of interest in red along with anti-MYO7A (hair cells) in blue, Lfng-EGFP in green and DAPI in white where applicable. Images to the right are the smFISH probe alone in grayscale. The inner hair cell (arrowhead) and outer hair cell regions (bracket) are indicated. *Myh9* = Myosin heavy chain 9 gene; *Cdkn1b* = Cyclin dependent kinase inhibitor 1b gene; *Lcp1* = Lymphocyte cytosolic protein 1 gene; *Notch2* = Notch receptor 2 gene; *Nlrp3* = NLR family pyrin domain containing 3 gene; *Slc2a3* = Solute carrier family 2 member 3 gene; TUBA1B = acetylated tubulin; MYO7A = myosin 7A protein; DAPI = 4’,6-diaminodino-2-phenylindole; Scale bars in all panels, 20 μm.

The heat map illustrates the top 800 differentially expressed genes between the two SC groups. Approximately 400 genes are localized to the upper half of the heat map while an approximately 400 additional genes that show generally higher expression in SC2 are localized in the lower half of the plot (Figure 2B). To examine the quality of the data set, we selected 6 genes, 2 known (*Myh9*, *Cdkn1b*) and 4 novel (*Lcp1*, *Notch2*, *Nlrp3*, *Slc2a3*), that showed expression within the single cell, to validate using smFISH and/or immunochtochemistry. *Myh9* and *Cdkn1b* have been reported to be expressed in most adult supporting cells (Mhatre et al. 2004; Löwenheim et al. 1999). Consistent with that, we observed uniform expression of both genes in SC1 and SC2 (Figure 2C). smFISH analysis also showed comparable levels of expression across the different supporting cells types (Figure 2D). *Lcp1* and *Notch2* showed similar broad patterns of expression within SC1 and SC2 although *Notch2* appeared to be expressed at lower levels in SC2 relative to SC1 (Figure 2E). smFISH results for *Lcp1* and *Notch2* were largely consistent with the scRNA-Seq results in that expression was observed in all supporting cells (Figure 2F). However, the relatively subtle difference in expression present in the single cell data was not obvious in any group of cells in the cross section. *Nlrp3* and to a lesser extent *Slc2a3* showed expression that appeared to be higher in SC1 versus SC2 (Figure 2E). smFISH results for Nlrp3 and Slc2a3 were inconsistent with scRNA-Seq results in that expression was observed in all supporting cells (Figure 2F). These conflicting results for Nlrp3 and Slc2a3 between scRNA-Seq and smFISH suggest that detection bias may account for decreased detection in one of the supporting cell clusters observed in scRNA-Seq. Detection bias or “dropout” refers to the event where a transcript is not detected in the sequencing data due to a failure to capture or amplify it (Haque et al. 2017). Overall, the validation of these markers in adult cochlear supporting cells points to the utility of this scRNA-Seq dataset for examining supporting cell-specific transcriptomes.

### Adult cochlear supporting cells can be categorized into two subpopulations

In order to gain a better overall perspective on the transcriptional diversity of adult cochlear supporting cells, examination of a larger group of differentially expressed genes was performed. A heatmap depicting the top 2460 transcripts expressed by the two identified clusters of supporting cells, SC1 and SC2 (Figure 3A) demonstrates some shared expression but also identifies a group of genes that distinguish SC2 from SC1. Two main clusters of genes are distinguished by the color bars on the vertical axis to the right of the heatmap with the red bar delineating 460 transcripts that demonstrate higher RNA expression in SC1 and the blue bar delineating 2000 transcripts that appear to exhibit higher RNA expression in SC2. Default parameters for significance included a minimum of 25% of cells expressing a given transcript and a fold change of at least 1.7. Despite the visually apparent similarity in SC1 and SC2 with respect to the 460 genes identified by the red bar, a composite expression plot of the SC1- and SC2-defining profiles demonstrates that these transcripts in composite appear to define two separate clusters of adult cochlear supporting cells, respectively (Figure 3B).

**Figure 3.**
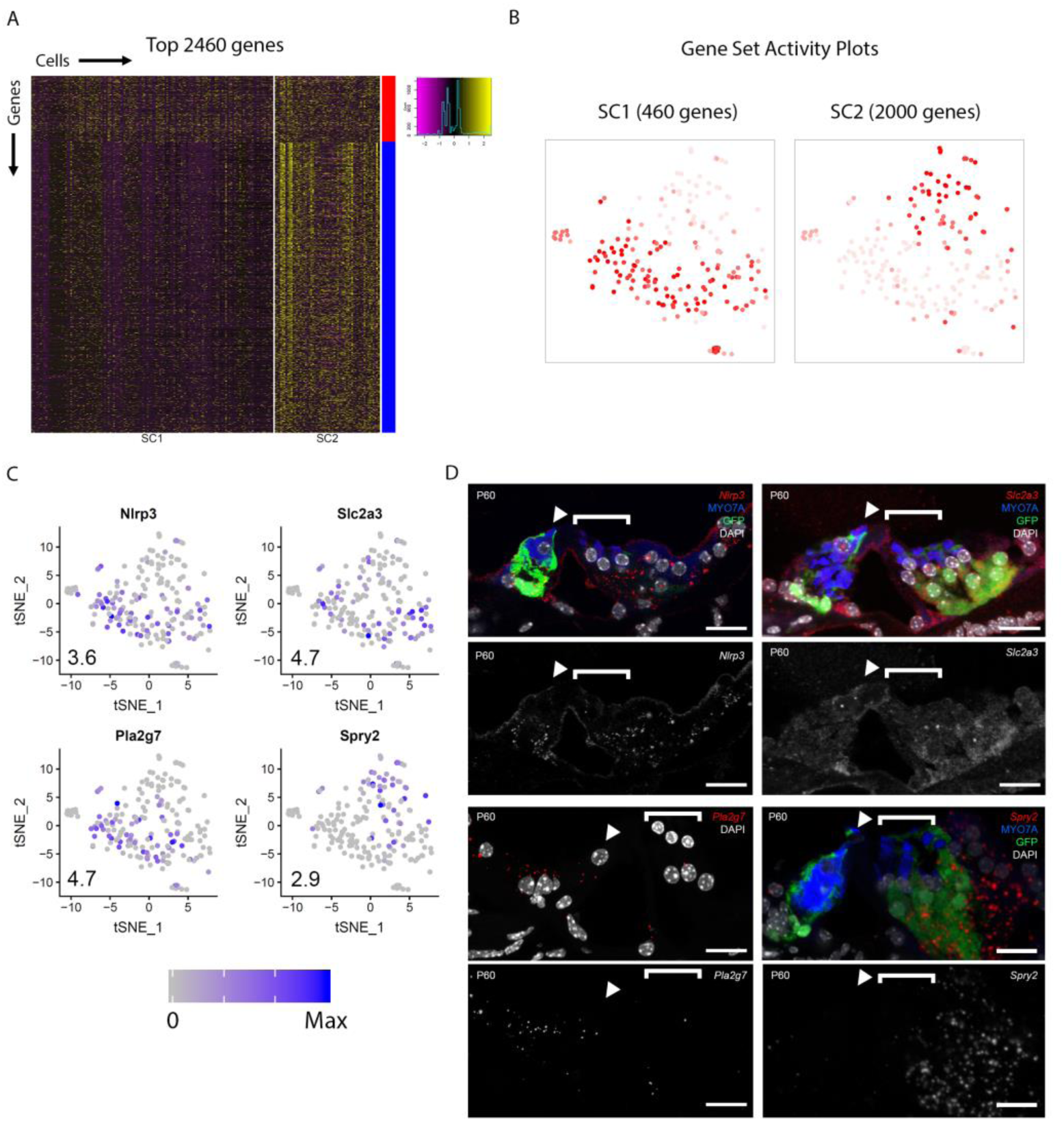
Adult cochlear supporting cells can be categorized into two subpopulations. **(A),** Heatmap depicting the top 2460 genes expressed by the two clusters of adult cochlear supporting cells (SC1, SC2). For SC1, only 460 genes met criteria for significance, while for SC2, 2000 genes that met criteria for significance were identified (see Methods). Cells are arrayed along the horizontal axis and genes are arrayed along the vertical axis. Two main clusters of genes are distinguished by the color bars on the vertical axis to the right of the heatmap with the red bar corresponding to SC2 and the blue bar corresponding to SC2. **(B),** Gene set activity plots demonstrate composite gene expression projected onto feature plots in SC1-defining (red bar, 460 genes) and SC2-defining (blue bar, 2000 genes). Despite the visually apparent presence of some SC1-defining genes in the SC2 cluster of supporting cells, SC1-defining genes as a composite appear to define the SC1 adult cochlear supporting cell subpopulation. **(C),** Feature plots of 1 known (*Tuba1b*) and 3 novel (*S100a6*, *Pla2g7*, *Spry2*) genes show differential expression between SC1 and SC2 supporting cell subpopulations. *S100a6* and *Pla2g7* correspond to SC1 (red gene cluster in Figure 3A) and *Tuba1b* and *Spry2* correspond to SC2 (blue gene cluster in Figure 3A). Numbers in the bottom left corner of the feature plots represent maximum expression level (Max) among adult cochlear supporting cells. Expression histogram is shown with blue indicating higher expression. **(D),** smFISH localization of RNA expression for these 4 candidate genes. Below each color image with the RNA probe color designated in italics is a grayscale single channel of the RNA probe. *S100a6* and *Pla2g7* demonstrate higher levels of transcript expression in the medial supporting cells (inner border, inner phalangeal cells) while *Spry2* demonstrates higher levels of transcripts in the lateral supporting cells (predominantly Deiters cells). While the immunological data in Figure 3D for an antibody the binds acetylated tubulin (TUBA1B) shows expression in pillar and Deiters cells, smFISH for *Tuba1b* in the grayscale single channel image shows low level expression in all SCs with a slightly greater level of Tuba1b transcripts in the lateral SCs (pillar and Deiters cells) compared to medial SCs (border and inner phalangeal cells). The region of the lateral SCs (pillar and Deiters cells) is identified by the dashed yellow line in the Tuba1b smFISH probe grayscale image. Hair cells are labeled when possible with MYO7A (blue), cell nuclei are labeled with DAPI (white) and Lfng-EGFP transgene expression (green) in supporting cells where applicable. The inner hair cell (arrowheads) and outer hair cell regions (bracket) are indicated. *S100a6* = S100 calcium-binding protein a6 gene; *Pla2g7* = Phospholipase A2 Group VII gene; *Tuba1b* = Tubulin alpha 1b gene; *Spry2* = Sprouty RTK signaling antagonist 2; TUBA1B = acetylated tubulin; MYO7A = myosin 7A protein; DAPI = 4’,6-diaminodino-2-phenylindole; Scale bar in all panels, 20 μm.

To examine whether the SC1 and SC2 might represent distinct supporting cell types, expression of four genes, including one known (*Tuba1b*) and three novel (*S100a6*, *Spry2*, *Pla2g7*) genes, which showed differential expression between SC1 and SC2 (Figure 3C) were examined in cross-sections. TUBA1B has previously been shown to be uniformly expressed in all adult SCs (Oesterle and Campbell 2009), however, the scRNAseq data indicated significant differential expression between the SC1 (low) and SC2 (high). The immunological data presented in Fig. 1D as well as the co-labeling with an antibody that binds acetylated tubulin (TUBA1B) in Figure 3D is consistent with that result showing higher expression of TUBA1B in lateral SCs (pillar cells and Deiters’ cells)(Figure 3D). While there is low level expression of *Tuba1b* transcripts in all SCs on smFISH, there appears to be greater levels of *Tuba1b* transcripts in the lateral SCs (pillar and Deiters cells) compared to medial SCs (border and inner phalangeal cells) with the lateral SCs demarcated by the yellow dashed lines in the grayscale single channel image for the *Tuba1b* RNA probe (Figure 3D). However, from the limited number of transcripts detected by smFISH it is difficult to assess whether any differences in transcriptional expression are present. In contrast with *Tuba1b*, *S100a6* showed higher expression in SC1 relative to SC2 (Figure 3C). Consistent with this result, smFISH indicated high levels of *S100a6* transcripts in medial SCs (border and inner phalangeal cells) and essentially no transcripts in lateral SCs. *Spry2* showed higher expression in SC2 relative to SC1 while *Pla2g7* showed higher expression in SC1 relative to SC2 (Figure 3C). Consistent with these results, smFISH indicated high levels of *Spry2* transcripts in lateral SCs (predominantly Deiters cells) and higher levels of *Pla2g7* transcripts in the medial SCs (border and inner phalangeal cells) relative to the lateral SCs (pillar and Deiters cells). Overall, these results confirm the expression of a subset of candidate genes in SCs and strongly suggest that SC1 and SC2 supporting cell subpopulations, represent medial and lateral SCs, respectively.

### Digital droplet PCR (ddPCR) and single-cell qPCR analyses of additional transcripts validate adult cochlear supporting cell scRNA-Seq

While validation using smFISH and/or immunohistochemistry provides valuable spatial information regarding transcriptional expression, these methods are labor intensive and low throughput. In our case, 2 cochlear supporting cell subpopulations were identified in the data and there was not necessarily a way to specifically isolate the exact combination of cells that created these subpopulations in our data from Lfng^EGFP^ adult mice. Therefore, we wanted to determine whether digital droplet PCR (ddPCR) and single-cell qPCR (sc-qPCR) could be used in combination as higher throughput methods for validating scRNA-Seq data. Specifically, the presence or absence of the genes of interest in FACS-purified GFP-positive adult cochlear supporting cells could be determined by ddPCR and the differential expression between two supporting cell subpopulations could be confirmed with sc-qPCR. For these reasons, we identified an additional group of 93 differentially expressed genes between SC1 and SC2 cochlear supporting cell subpopulations that were selected from the top 800 differentially expressed genes. Some genes were chosen due to preexisting knowledge about cochlear supporting cells while the remainder of the candidate genes were chosen at random. Using a subset of these genes, we confirmed the presence of their transcripts in populations of FACS-purified adult cochlear supporting cells using ddPCR where presence was measured in number of copies per 2,000 cells (Figure 4A). Expression was present and quantifiable in each of the genes from this selected group of genes. In order to validate the differential expression of genes between the two cochlear subpopulations (SC1 and SC2), we performed sc-qPCR on 170 FACS-purified GFP-positive cochlear supporting cells from the Lfng^EGFP^ adult mouse cochlea using this additional group of differentially expressed genes. Principal component analysis (PCA) corroborates unbiased clustering of single cell transcriptomes into 2 clusters on the basis of these genes (Figure 4B). Expression of these genes is depicted in the heatmap (Figure 4C). Violin plots show expression level of each of these genes as determined by sc-qPCR (Figure 4D). These data demonstrate the potential utility of a combinatorial approach to scRNA-Seq validation utilizing ddPCR to screen for the presence or absence of transcripts in a mixed population and then utilizing sc-qPCR to validate differential expression observations.

**Figure 4.**
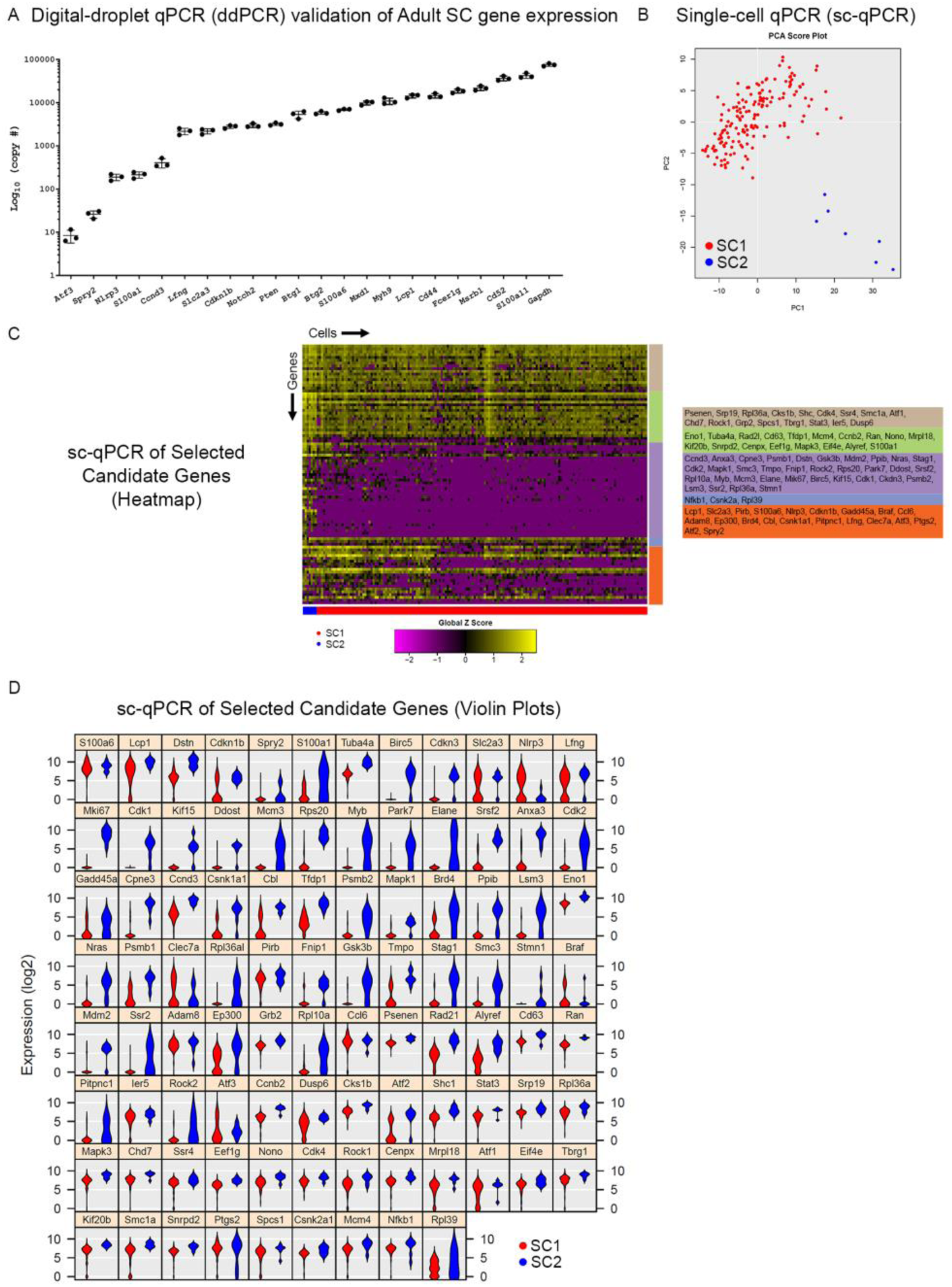
Digital droplet PCR (ddPCR) and single-cell qPCR analyses of additional transcripts validate adult cochlear supporting cell scRNA-Seq. **(A),** Digital droplet PCR (ddPCR) quantification of candidate genes in FACS-purified P60 *Lfng^EGFP^*-positive cochlear supporting cells. Absolute quantitation measured in number of transcript copies detected are plotted on the log base 10 scale on the vertical axis and genes of interest are on the horizontal axis. All candidate genes were detected indicating expression in FACS-purified adult cochlear supporting cells. **(B),** Principal component analysis (PCA) of single cell qPCR (sc-qPCR) of 170 FACS-purified P60 Lfng*^EGFP^*-positive cochlear supporting cells corroborates unbiased clustering of single cell transcriptomes into 2 clusters (SC1 in red, SC2 in blue). **(C),** Heatmap of sc-qPCR results with 170 adult cochlear supporting cells (along horizontal axis) and 93 candidate genes (along vertical axis). Candidate gene clusters as determined by hierarchical clustering are noted as the colored bars along the vertical axis of the heatmap. Candidate genes making up the gene clusters are noted to the right of the heatmap in the corresponding colored boxes. As denoted by the expression histogram, the higher the expression, the more yellow the box corresponding to the gene in a given cell. **(D),** Violin plot display of sc-qPCR results demonstrates candidate gene expression levels in adult cochlear supporting cells.

### Supporting cell-specific genes are also expressed in human temporal bones from patients with or without intact hair cells

To determine whether SC markers identified in mice are also expressed in human SCs, we localized the expression of two novel SC markers, S100A6 and LCP1, in organs of Corti from human temporal bone specimens. Prior to attempting immunolocalization in human specimens, we validated antibodies for S100A6 and LCP1 in adult mouse organ of Corti, as smFISH has not been previously demonstrated to work in archival temporal bone specimens. S100A6 protein is expressed in predominantly medial SCs (border and inner phalangeal cells) with some expression noted in the Hensen’s cells in the adult mouse as seen on mid-modiolar cross-sections of the adult organ of Corti (Figure 5A-A’). Grayscale single channel images of S100A6 and LCP1 protein expression are shown in Figures 5A’ and 5B’, respectively. S100A6 protein expression in Hensen’s cells appears to be intermediate to expression in inner border/inner phalangeal cells and Deiters cells, respectively. LCP1 protein was localized predominantly to the pillar cells (Figure 5B-B’). This differs results of smFISH in Figure 2 where the distribution of *Lcp1* RNA transcripts is observed in all cochlear supporting cells (Figure 2F). This observed incongruence between *Lcp1* RNA and LCP1 protein could be due to undefined post-transcriptional processing mechanisms, differences in the half-life of the LCP1 protein in vivo, or imperfect binding of the antibody to the LCP1 protein (Greenbaum et al. 2003).

**Figure 5.**
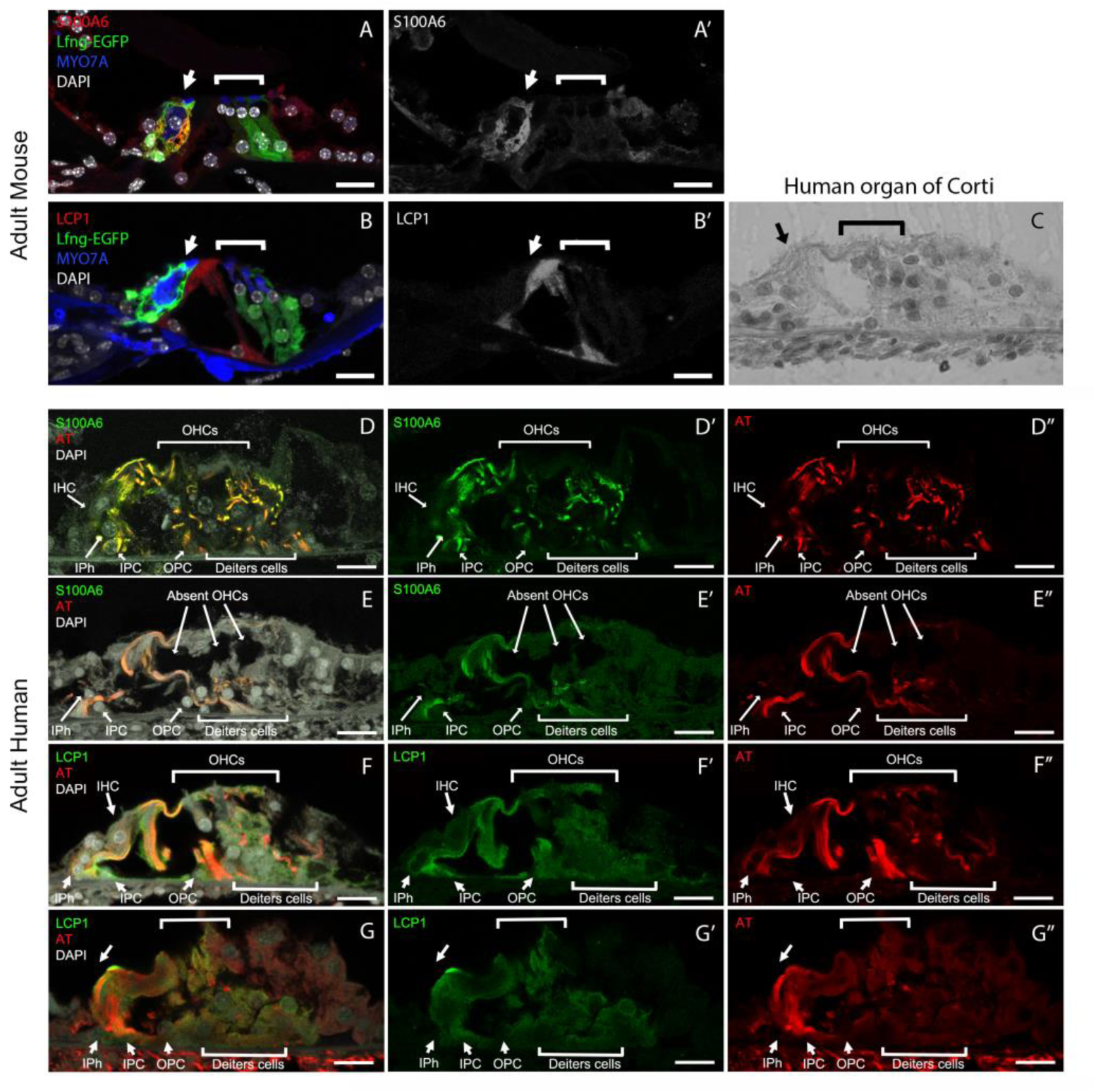
Supporting cell-specific genes are also expressed in human temporal bones from patients with or without intact hair cells. **(A-A’),** S100A6 protein expression in a representative mid-modiolar cross section of P60 Lfng^EGFP^ organ of Corti. S100A6 protein (red) is localized to GFP-positive supporting cells (A) and grayscale single channel image demonstrates S100A6 protein expression (A’). (**B-B’),** LCP1 protein expression in a representative mid-modiolar cross section of P60 Lfng^EGFP^ organ of Corti. LCP1 protein (red) is localized to GFP-positive supporting cells (B) and grayscale single channel image demonstrates LCP1 protein expression primarily in pillar cells (B’). Myosin 7A (MYO7A) and DAPI co-labeling for hair cells and nuclei, respectively. (**C),** Brightfield image of a human organ of Corti is shown with an arrow pointing to the region of the inner hair cell and a bracket outlining the outer hair cell region. (**D-D”),** Representative immunofluorescence of celloidin embedded human organ of Corti sections from a patient with normal hearing demonstrate S100A6 expression (green) that overlaps with acetylated tubulin (AT), a known adult cochlear supporting cell marker (red) (C). Single channel images for S100A6 in green (C’) and acetylated tubulin (AT) in red (C”) are shown. Arrow and bracket point to the inner hair cell (IHC) and outer hair cell (OHC) region, respectively. (**E-E”),** S100A6 expression in human organ of Corti from a patient with age-related hearing loss and hair cell loss. Note that S100A6 expression persists in the absence of hair cells. Representative immunofluorescence demonstrates S100A6 (green) and AT (red) (D) with single channel images for S100A6 protein (D’) and AT (D”). Arrow and bracket point to IHC and OHC regions, respectively and are notable for a lack of these cell types in the section. (**F-F”),** Representative immunofluorescence of celloidin embedded human organ of Corti sections from a patient with relatively normal hearing demonstrate LCP1 expression (green) overlaps with AT (red) (E). Single channel images for LCP1 in green (E’) and AT in red (E”) are shown. The location of inner and outer hair cells are marked by the arrow and bracket, respectively. (**G-G’’),** LCP1 expression in human organ of Corti from a patient with age-related hearing loss and hair cell loss. Arrow and bracket point to IHC and OHC regions, respectively and are notable for a lack of these cell types in the section. Note that LCP1 expression persists in the absence of hair cells. S100a6 = S100 calcium-binding protein a6; LCP1 = lymphocyte cytosolic protein 1; AT = acetylated tubulin; MYO7A = myosin 7A protein; DAPI = 4’,6-diaminodino-2-phenylindole; IHC = inner hair cell; OHC = outer hair cell; IPh = inner phalangeal cell; IPC = inner pillar cell; OPC = outer pillar cell. Scale bar in all panels, 20 μm.

An antibody against acetylated tubulin was utilized to label adult human cochlear supporting cells and confirm cell type-specific expression in human samples. A brightfield image of a human organ of Corti is shown in Figure 5C with an arrow pointing to the region of the inner hair cell and a bracket outlining the outer hair cell region. S100A6 and LCP1 expression are demonstrated in adult human cochlear supporting cells and co-localize with acetylated tubulin, a known adult supporting cell marker (Figure 5D-D”, 5F-F”). Unlike S100A6 protein in mouse, S100A6 protein in humans appears to be expressed by all cochlear supporting cells. Between mouse and human, LCP1 protein expression appears to correlate with expression noted in the pillar cells. Furthermore, these proteins are expressed in the human organ of Corti in the absence of hair cells demonstrating their potential relevance to future therapies targeting supporting cells in the adult human inner ear (Figure 5E-E”, 5G-G”). Unlike acetylated tubulin, which is expressed in the human stria vascularis, LCP1 is not expressed in the stria vascularis (Supplemental Figure S7). These data demonstrate that S100A6 and LCP1 protein are expressed by adult human cochlear SCs. Overall, our data suggest that other supporting cell markers identified in the single cell transcriptome dataset of adult mouse cochlear SCs could be reliable markers for adult human SCs.

### Cell cycle phase analysis reveals that the SC2 adult supporting cell subpopulation expresses transcripts associated with S phase and G2/M phase canonical markers

Focusing on mechanisms that might potentially facilitate mitotic regeneration in post-mitotic adult cochlear supporting cells, we utilized a recently-developed methodology to computationally assign cell cycle phase (G0/G1, S, or G2/M) based on cell cycle-specific gene expression to single cells on the basis of their single-cell transcriptomes (Scialdone et al. 2015; Nestorowa et al. 2016) (http://satijalab.org/seurat/cell_cycle_vignette.html). PCA analysis suggests that the SC2 supporting cell subpopulation which likely represents the lateral supporting cells (pillar and Deiters cells) (Figure 3D) express transcripts that are consistent with S phase or G2/M phase while SC1 cochlear supporting cells do not (Figure 6A-B and Supplemental Figure S8-9). Single cell qPCR in Figure 6C validates these findings with select S phase (*Mcm4*) and G2/M (*Birc5, Cdk1, Mki67*) phase transcripts showing either predominant expression (*Birc5*, *Cdk1*, *Mki67*) or mild enrichment in transcript expression (*Mcm4*) in SC2 (Figure 6C). These data validate baseline expression levels of these selected S phase- and G2/M phase-associated transcripts. The validation lends support to the idea that the scRNA-Seq data could be utilized to assess baseline expression levels of S phase- and G2/M phase-associated genes, which could be modulated in the future in order to facilitate mitotic regeneration.

**Figure 6.**
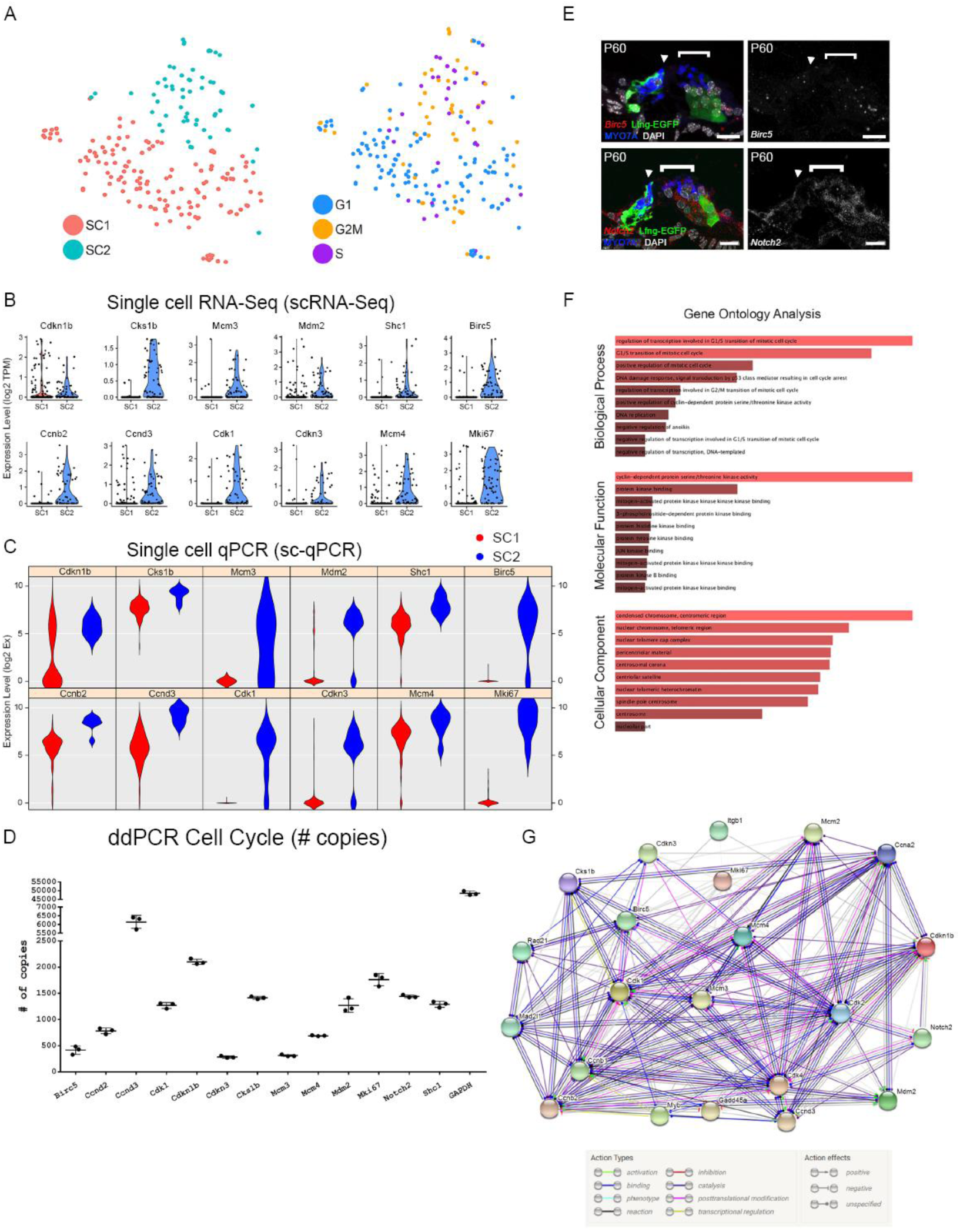
Cell cycle analysis of adult cochlear supporting cells. (**A),** Analysis of cell cycle phase among FACS-purified adult cochlear supporting cells reveals that the SC2 cluster of adult cochlear supporting cells express predominantly S phase and G2/M phase canonical markers compared to SC1 adult cochlear supporting cell cluster. tSNE plot with the clustered cell types is shown to the left with an accompanying tSNE plot demonstrating cells clustered by cell cycle phase to the right. (**B),** Single cell RNA-Seq expression of G2/M and S phase-specific markers. Violin plots for a select group of G2/M and S phase-specific cell cycle-related genes demonstrate predominant expression in the SC2 cluster of adult cochlear supporting cells. For violin plots, expression level (log2 TPM) is displayed on the vertical axis and cell cluster is displayed on the horizontal axis. (**C),** Single-cell qPCR of FACS-purified adult cochlear supporting cells for G2/M and S phase cell cycle-related markers confirms gene expression in adult cochlear supporting cells. Violin plots are redisplayed from Fig. 4D for ease of comparison to scRNA-Seq results above. (**D),** Digital droplet PCR (ddPCR) quantifies the presence of the select group of cell cycle-related genes validated by sc-qPCR in FACS-purified P60 Lfng*^EGFP^*-positive cochlear supporting cell populations. (**E),** Cell cycle gene expression is demonstrated in mid-modiolar sections of P60 mouse organ of Corti. RNA probe (in red) with accompanying immunohistochemistry is shown in image to the left and grayscale single channel image is shown in image to the right. smFISH probe for *Birc5* (red dots) are shown to overlap with adult cochlear supporting cells (GFP in green). Hair cells are labeled with MYO7A (blue) and nuclei are labeled with DAPI (white). Location of inner hair cell (arrowhead) and region of outer hair cells (bracket) are denoted. (**F),** Gene ontology (GO) analysis of cell cycle-related genes expressed by FACS-purified GFP-positive cells from P60 Lfng*^EGFP^* cochlea suggests that these cells may be maintained in a non-proliferative state by a repressive network of genes. All cell cycle genes expressed by adult supporting cells from the dataset, regardless of which cluster of adult cochlear supporting cells expressed these genes, were used as the starting input in Enrichr. GO biological process analysis suggests that genes involved in the G1/S transition of the mitotic cell cycle are prominent in adult cochlear supporting cells. GO molecular function and cellular component analysis point to cyclin-dependent protein serine/threonine kinase activity and cellular components associated with condensed chromatin at the centromere, respectively. The color of the bar corresponds to the combined score which is calculated by taking the log value of the p-value from the Fisher exact test and multiplying this value by the z-score of the deviation from the expected rank. The longer and lighter colored bars indicate that the term is more significant. (**G),** Use of the STRING database to perform protein-protein interaction analysis identifies a set of interactions that may be related to the persistence of the post-mitotic state in adult cochlear supporting cells. The STRING plot demonstrates the action types and action effects as noted in the accompanying legend. Scale bar in all panels, 20 µm.

### Adult cochlear supporting cells continue to express some cell cycle genes expressed by neonatal Lgr5+ inner ear progenitors

One of the reasons for the interest in adult supporting cells is the thought that they might serve as a potential stem cell reservoir for regenerating hair cells in the organ of Corti. Lgr5+ neonatal cochlear cells are thought to be potential progenitors for hair cells and a potential stem cell reservoir within the organ of Corti (F. Shi, Hu, and Edge 2013; T. Wang et al. 2015; McLean et al. 2016; Cheng et al. 2017). For these reasons, similarities between Lgr5+ neonatal cochlear supporting cells and adult cochlear supporting cells gene expression were examined. Cheng and colleagues surveyed genes expressed by Lgr5+ neonatal cochlear supporting cells and reported expression of genes implicated in cell cycle, Notch, EGF and Wnt signaling pathways (Cheng et al. 2017). To examine whether some of these same genes might continue to be expressed in adult supporting cells, and in particular in the SC2 group, violin plots for a subset of these cell cycle genes were generated (Supplemental Figure S10). Using ddPCR, we confirmed the presence of the transcripts for a selected group of these cell cycle genes in FACS-purified GFP-positive adult cochlear supporting cells (Figure 6B). Differential expression of these cell cycle genes was then validated by sc-qPCR (Figure 4D, Figure 6C). Results from the sc-qPCR from Figure 4D are redisplayed in Figure 6C to show correlation between scRNA-Seq violin plots (Figure 6D) and the sc-qPCR violin plots (Figure 6C). Transcripts identified as being enriched in Lgr5-positive neonatal supporting cells and enriched in adult cochlear supporting cells by both scRNA-Seq and sc-qPCR include *Cdkn1b, Cks1b, Mcm3, Mdm2, and Shc3*. Transcripts identified as being enriched in Lgr5-negative neonatal supporting cells and enriched in adult cochlea supporting cells by both scRNA-Seq and sc-qPCR include *Birc5*, *Ccnb2*, *Ccnd3*, *Cdk1*, *Cdkn3*, *Mcm4*, and *Mki67*. *Cdkn1b, Mcm3*, *Mdm2*, *Birc5*, *Cdk1*, *Cdkn3*, and *Mki67* exhibit relatively good agreement between supporting cell clusters in both scRNA-Seq and sc-qPCR. However, transcript expression, as determined by sc-qPCR, for *Cks1b, Shc1, Ccnb2, Ccnd3*, and *Mcm4* appears to contradict the results of scRNA-Seq in that expression is noted in the SC1 supporting cell cluster. Reasons for this discrepancy between scRNA-Seq and sc-qPCR include the likelihood that PCR may be more sensitive at detecting transcripts given its targeted nature while scRNA-Seq must contend with potential inefficiencies in reverse transcription and amplification. smFISH validation of select cell cycle genes identified by Nestorowa and colleagues (2016) and Cheng and colleagues (2017) (*Notch2*, *Birc5*) is also performed in adult mouse organ of Corti and their mRNA expression overlaps with adult GFP-positive cochlear supporting cells (Figure 2F and 6E) (Nestorowa et al. 2016; Cheng et al. 2017). A description of these candidate cell cycle genes whose transcripts are validated in adult cochlear supporting cells is noted in Supplemental Table S6. Shared gene expression of adult cochlear supporting cells with Lgr5+ neonatal cochlear supporting cells suggest that adult supporting cells may still possess latent abilities to serve as a stem cell reservoir.

### Adult cochlear supporting cells may be maintained in a non-proliferative state

Finally, to examine other pathways that are active in adult supporting cells, a gene ontology biological process analysis was performed. Results suggest that genes involved in the G1/S transition of the mitotic cell cycle are more prominently expressed in adult cochlear supporting cells by comparison with Lgr5+ neonatal cochlear supporting cells (Figure 6F). These genes may function to prevent adult supporting cells from re-entering or moving forward successfully through the phases of the cell cycle. In addition, gene ontology molecular function and cellular component analyses group these cell cycle genes to cyclin-dependent protein serine/threonine kinase activity and cellular components associated with condensed chromatin at the centromere, respectively, suggesting that these pathways could be activated in adult cochlear supporting cells and contribute to the maintenance of quiescence (Figure 6F). Use of the STRING database to perform protein-protein interaction analysis identifies a set of interactions that may be related to the persistence of the post-mitotic state in adult cochlear supporting cells (Figure 6G). Overall, these analyses suggest that these data could be utilized as a resource to explore mechanisms related to the maintenance of supporting cell quiescence.

## Discussion

One of the major causes of sensorineural hearing loss in adults, a significant and irreversible health problem, is the loss of sensory hair cells. While adult cochlear supporting cells may represent a potential source of replacement hair cells, hearing restoration will likely require regenerating the proper complement of cell types including any supporting cells that are converted to hair cells and restoration of the architecture of the organ of Corti, (Jahan et al. 2015). The exact temporal expression sequence and level of gene expression required for the creation of both mature hair cells and supporting cells is yet to be elucidated. Furthermore, the adult organ of Corti appears to contain barriers to regeneration that remain incompletely defined. Improved understanding of the final mature state of cell types in the organ of Corti may contribute to future attempts at hearing restoration through regenerative approaches. These data provide gene expression levels for two groups of cochlear supporting cells that are targets for induction of regenerated hair cells. Many of the genes we have identified in these supporting cells have not been previously characterized in the inner ear and may be examined as possible targets for the development of supporting cell-specific gene expression or as candidates for induction of changes in cell fate or mitotic state (Supplemental Table S2).

### Adult cochlear supporting cells are transcriptionally distinct from perinatal cochlear supporting cells

While previous studies have focused on identifying the genetic cascade necessary to convert perinatal cochlear supporting cells into hair cells (Liu et al. 2012; Kelly et al. 2012; Walters et al. 2017; Costa et al. 2015), it seems possible that changes in the transcriptome and/or epigenetic state of adult supporting cells may necessitate activation of different pathways. Consistent with this hypothesis, the results presented here demonstrate that the transcriptomes of adult cochlear supporting cells are markedly different from P1 cochlear supporting cells (Burns et al. 2015)(GEO Accession ID: GSE71982; Figure 1). Biological process gene ontology analysis reveals that P1 cochlear supporting cells demonstrate enrichment for genes involved in positive regulation of Wnt signaling (GO:0090263, GO:0030177), regulation of stem cell differentiation (GO:2000736), and mitotic cell cycle phase transition (GO:1901990), all of which could play a role in a regenerative response. In contrast, adult cochlear supporting cells demonstrate enrichment for genes involved in regulation of MAP kinase activity (GO:0043406, GO:0043405, GO:0000187), regulation of JNK cascade (GO:0046328), activation of protein kinase activity (GO:0032147), and integrin-mediated signaling pathway (GO:0007229). Work by others has suggested that modulation of MAP kinase signaling, JNK signaling and protein kinase signaling may facilitate S-phase entry and proliferation (Montcouquiol and Corwin 2001). These authors suggest that a balance between MAP kinase signaling and JNK signaling may determine whether cells proliferate or die by apoptosis. Work in zebrafish demonstrates that inhibition of JNK signaling leads to suppression of hair cell regeneration apparently by preventing proliferation, suggesting that JNK signaling plays a role in proliferation in inner ear sensory epithelia (He et al. 2016). However, Montcouquiol and Corwin suggest that activation of these signaling pathways (MAP kinase, JNK, protein kinase activity) may not be sufficient for proliferation. Davies and colleagues have also suggested that changes in the cell-extracellular matrix interactions as cells mature in the cochlea, notably changes in integrin expression, may maintain these cells in their postmitotic quiescent state (Davies, Magnus, and Corwin 2007). These observations combined with the absence of enrichment for genes involved in Wnt signaling and mitotic cell cycle transition may highlight some key differences between adult and perinatal cochlear supporting cells. Overall, these observations point to a need to better understand what distinguishes adult cochlear supporting cells from perinatal cochlear supporting cells (White et al. 2006). While it may seem obvious that adult cochlear supporting cells are transcriptionally distinct, much of the current work into elucidating the developmental transitions necessary for the production of new hair cells and supporting cells presumes that these same developmental transitions can be applied to adult cochlear supporting cells *in vivo*.

### Single-cell RNA-Seq identifies adult supporting cell gene expression

In order to demonstrate the utility of this dataset, we utilized smFISH to both confirm and localize transcript expression to adult cochlear supporting cells. smFISH validation of both known (*Myh9, Cdkn1b*) and novel (*Lcp1, Notch2*) genes shows relatively good concordance with scRNA-Seq data. While scRNA-Seq results for *Nlrp3* and *Slc2a3* show higher expression in SC1 compared to SC2, smFISH demonstrates presence across all supporting cells. Reasons for discrepancies between scRNA-Seq and sc-qPCR include the likelihood that PCR may be more sensitive at detecting transcripts given its targeted nature, while scRNA-Seq can show decreased sensitivity for particular transcripts because of inefficiencies in reverse transcription and amplification.

### Adult cochlear supporting cells can be categorized into two subpopulations

Our data identified two transcriptionally distinct supgroups of adult supporting cells (SC1 and SC2). smFISH localization of *S100a6* and *Pla2g7*, markers of SC1, was largely restricted to medial SCs (border and inner phalangeal cells), while *Tuba1b* and *Spry2,* markers of SC2, were localized to lateral SCs (pillar and Deiters cells). More importantly, the composite expression of each of these two identified clusters of genes define these two groups as seen on the gene set activity plots (Figure 3B) and support the potential relevance of combinatorial gene expression in identifying subpopulations (Patel et al. 2014). These results are consistent with single cell analysis of the developing cochlear duct which suggest that the organ of Corti is derived from two transcriptionally distinct regions, medial and lateral, with medial giving rise to inner hair cells and associated supporting cells and lateral giving rise to outer hair cells and associated supporting cells (Kolla and Kelley, personal communication). Since these regional distinctions could also be indicative of lineage restrictions, the identification of markers that define each domain could be utilized to create gene therapy targeting vectors specific for medial or lateral supporting cell types (Stone and Cotanche 2007; Cox et al. 2014).

### Population qPCR and single-cell qPCR validation of additional gene candidates from adult single cell transcriptomes

Recently, single-cell RNA-sequencing through commercial droplet-based platforms has achieved higher throughput in comparison to microfluidics-based platforms, such asthe Fluidigm C1 platform utilized in this study. However, because the Fluidigm system does not rely on molecular barcodes to discriminate cell of origin, ddPCR and single-cell qPCR can be used to examine expression of specific transcripts within each data set, with a higher level of sensitivity by comparison with scRNA-seq (Figure 4, Figure 6). The results of our analysis of supporting cells using ddPCR and/or sc-qPCR yielded results that were largely consistent with the scRNAseq results (Figure 4, Figure 6). However, some differences were also observed. For instance, while *S100a6* transcript expression is increased in SC1 versus SC2, a result that was observed by smFIHS as well, single cell qPCR (sc-qPCR) for *S100a6* demonstrates relatively equivalent levels of gene expression. This result could be a result of sampling bias, as only 7 SC2 cells were collected for the sc-qPCR data set or might reflect decreased amplification efficiency for this primer set. Future efforts to increase the number of SC2 cells may address some of the discrepancies in between the different methods.

### Adult mouse cochlear supporting cell-specific genes are expressed in human inner ears

An important consideration in any biomedically-related study using an animal model is the applicability to humans. For this study, we examined the expression of two candidate adult mouse SC genes in human cochlear SCs. S100A6 and LCP1 demonstrated slightly different patterns of expression in humans versus mice. Specifically, S100A6 protein appears to be expressed in most human supporting cells (Figure 5D-E”) unlike the mouse where S100A6 protein expression is more highly expressed in medial SCs (Figure 5A-A’). In addition to transcriptional differences between mouse and humans, S100A6 is known to be secreted and taken up by other cells, which may explain this apparent disparity in protein expression between mouse and human SCs (Jurewicz, Wyroba, and Filipek 2018). In contrast to the observed broad expression *Lcp1* RNA across all supporting cells in the mouse (Figure 2F), LCP1 protein expression in the human organ of Corti appears to be confined to the pillar cells (Figure 5B-B’). Potential reasons for this disparity include insufficiently defined post-transcriptional mechanisms involved in converting mRNA into protein, differing in vivo half-lives of protein, and the existence of a significant amount of error and noise in both protein and mRNA experiments (Greenbaum et al. 2003). Greenbaum and colleagues also suggest that changes in the rate of protein and mRNA synthesis as well as protein turnover may contribute to this disparity. LCP1 protein expression in humans is consistent with observed protein expression in the mouse (Figure 5F-G”). More importantly, the expression patterns of both proteins are preserved in the face of human hair cell loss, which is a crucial consideration in terms of the identification of potential drivers for induction of a regenerative paradigm in SCs (Figure 5E-E”, G-G”). These data also point to the utility of human temporal bone repositories as a resource to validate and confirm candidates for hair cell regeneration attempts in humans.

### A potential role for cell cycle regulation in adult cochlear supporting cells

A pathway analysis for the SC2 subpopulation indicated expression of canonical S and G2/M phase genes and similarities to Lgr5-expressing cells isolated from neonatal cochlea (Nestorowa et al. 2016; Cheng et al. 2017). These Lgr5-expressing cells in the neonatal cochlea possess a latent potential to proliferate and to convert into hair cells (Cox et al. 2014; Stone and Cotanche 2007). This mitotic regeneration has been explored in supporting cells as a potential avenue for hair cell regeneration with Wnt activation and Notch signaling playing roles in supporting cell conversion into hair cells (Ni et al. 2016; McGovern et al. 2018, 2019). Consistent with these observations, protein interaction analyses suggest potentially relevant interactions between Notch signaling and Wnt signaling pathways and cell cycle control, specifically *Cdkn1b* (Figure 6G). Work by others suggests that these signaling pathways may be relevant to achieving hair cell regeneration in the adult inner ear (Hori et al. 2007; Kuo et al. 2015; Walters et al. 2017).

While overall biological process gene ontology analysis suggests involvement in the G1/S transition of the mitotic cell cycle, sub-analyses categorize these genes into three main categories: positive regulation of transcription from RNA polymerase II promoter, G1/S transition of mitotic cell cycle, and negative regulation of DNA endoreduplication. A supplemental table of these genes and references regarding their possible functional roles in cell cycle is provided (Supplemental Table S6).

With regards to the first gene ontology term (positive regulation of transcription from RNA polymerase II promoter), in other systems these genes are involved in either directly promoting proliferation or facilitating proliferation by overcoming cellular defense mechanisms to cell cycle reentry (see Supplemental Table S6 for full set of references). Overcoming cellular defense mechanisms activated in response to cell cycle reentry may be critical to achieving adult supporting cell proliferation (Ping Chen et al. 2003).

The genes categorized under the second gene ontology term (G1/S transition of mitotic cell cycle) are related to the maintenance of quiescence in adult cochlear SCs (Malgrange et al. 2003; P Chen and Segil 1999; Laine et al. 2010; Oesterle et al. 2011). Specifically, inhibition of cyclin-dependent kinases (CDKs) and cyclin-dependent kinase inhibitors (CKIs), some of which are expressed by adult cochlear SCs, have been implicated in the generation of supernumerary neonatal hair cells and supporting cells (Malgrange et al. 2003; P Chen and Segil 1999; Laine et al. 2010; Oesterle et al. 2011). Transcript expression levels for these and other cell cycle-related genes potentially represent thresholds for cell cycle re-entry. Expression of these genes may be indicative of a poised state with modulation of these genes by subtle overexpression or inhibition around the baseline potentially leading to stimulation of cellular proliferation (Hu et al. 2016). This poised state has also been suggested by the discovery of bivalent chromatin structure in perinatal supporting cells (Hu et al. 2016; Stojanova, Kwan, and Segil 2015) and may exist in adult supporting cells, albeit with additional barriers to regeneration in place.

Finally, with these additional barriers in mind, the genes categorized under negative regulation of DNA endoreduplication are related to facilitating but not necessarily initiating cell proliferation (see Supplemental Table S6 for full set of references). Endoreduplication refers to re-replication of DNA within a single cell, likely representing a collection of compensatory mechanisms for maintaining viability of cells that are unable to progress completely through the cell cycle during the course of regeneration (Denchi, Celli, and De Lange 2006). Modulation of this group of genes may facilitate the creation of a genomic environment that is more conducive to cell proliferation, which under normal circumstances would be prevented/inhibited by several other factors. For example, while p53 inactivation through an inducible conditional p53 knockout mouse does not result in adult supporting cell proliferation or transdifferentiation into hair cells in mice, the loss of p53 may create an environment where supporting cell proliferation is possible given appropriate but currently undefined conditions (Laos et al. 2017).

Despite these analyses, it is possible that inducing adult supporting cell proliferation will require expression of cell cycle-related genes that are not normally transcribed by adult cochlear SCs. Furthermore, despite observations about the connections between Wnt and Notch signaling pathways and cell cycle control, others have noted that the presence of Wnt signaling components in adult mouse cochlea in the face of an inability for supporting cell mitosis or trans-differentiation argues for the existence of epigenetic mechanisms preventing supporting cells from utilizing these machinery (Geng et al. 2016; Samarajeewa, Jacques, and Dabdoub 2019). Interactions between existing cell cycle gene expression and epigenetic machinery may maintain quiescence in adult cochlear supporting cells and remain incompletely elucidated. Nonetheless, the results presented here indicate the expression of a wide network of cell cycle genes in adult cochlear SCs, suggesting that a better understanding of the roles of these networks in maintaining adult supporting cell quiescence could provide valuable insights regarding pathways to hair cell regeneration. Future studies will be required to test the relevance of modulating cell cycle gene expression or modulating interactions between cell cycle genes and epigenetic machinery to achieving hair cell regeneration in the adult cochlea. These data suggest that there are likely multiple mechanisms that may be involved in maintaining the latent potential of cochlear supporting cells to re-enter the cell cycle and to transdifferentiate into hair cells. Understanding the potentially different avenues of overcoming this poised state in adult cochlear supporting cells may lead to more effective hair cell regeneration in the future.

## Conclusions

In summary, we have used several different approaches, including sc-RNAseq, ddPCR and sc-qPCR, to characterize and validate the transcriptional profiles of cochlear supporting cells from adult mice. Our results demonstrate significant changes in gene expression between perinatal and adult supporting cells. To determine whether these results may be relevant to efforts to induce regeneration in humans, we examined expression of two of these genes in human temporal bones; which demonstrated patterns of expression in supporting cells that were comparable to those of adult mice. Finally, analysis of gene expression in adult supporting cells indicates strong expression of pathways related to regulation of the cell cycle, suggesting that targeting of these pathways could help to force cells out of quiescence. These results provide insights that could be relevant to the development of treatments to induce hair cell regeneration, which will, most likely, include the trans-differentiation of supporting cells into new hair cells, leading to a deficit in supporting cells. If this is the case, controlled creation of new supporting cells, through cell cycle re-entry, may be a crucial step in the restoration of a functional cochlea.

**Table 1.**
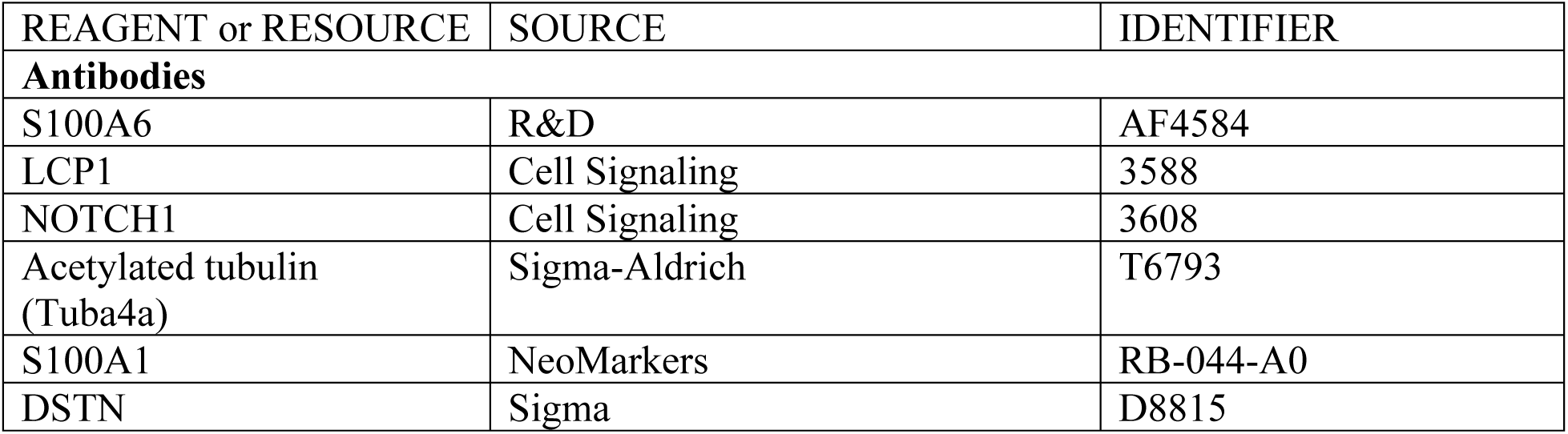
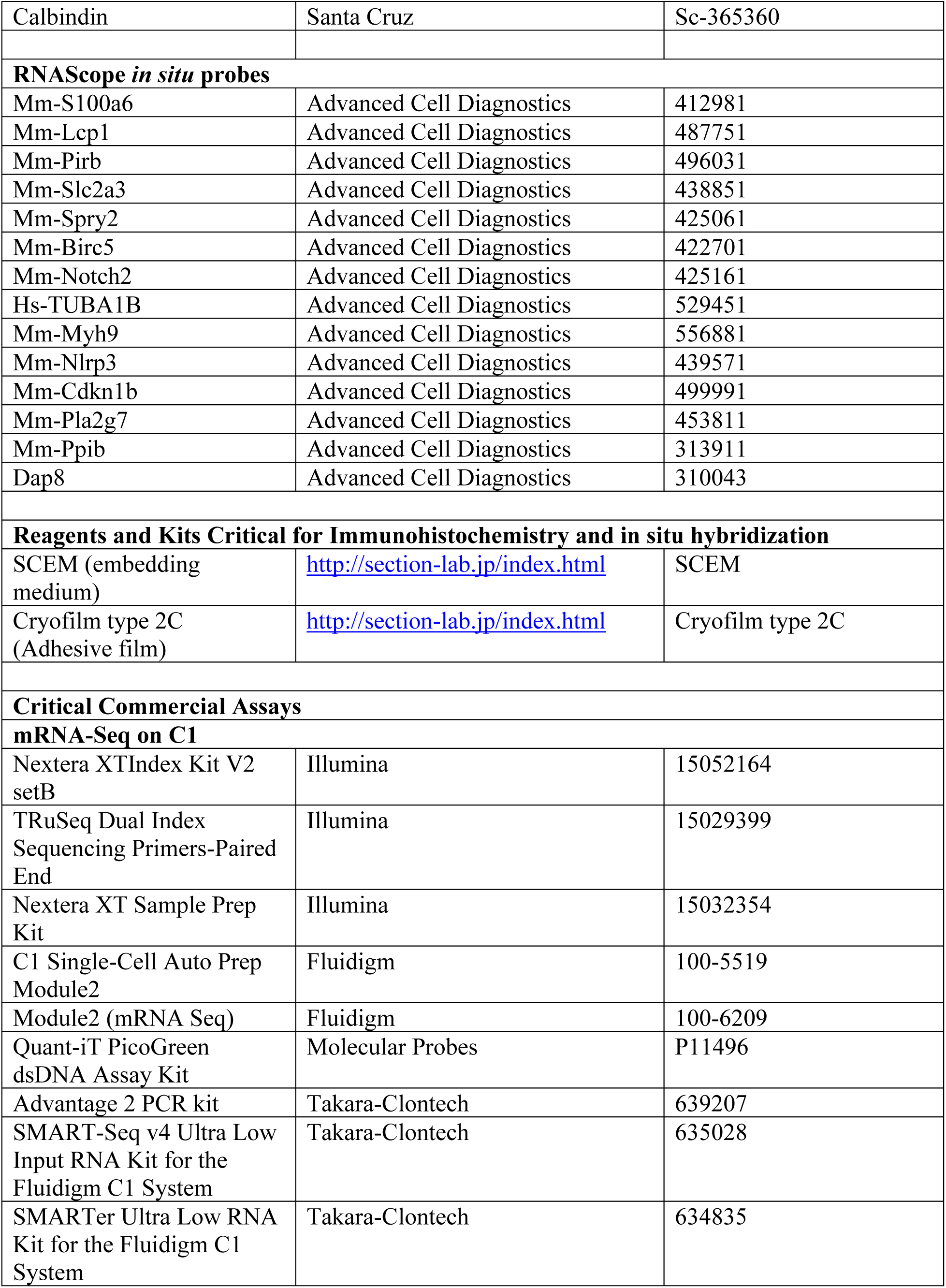
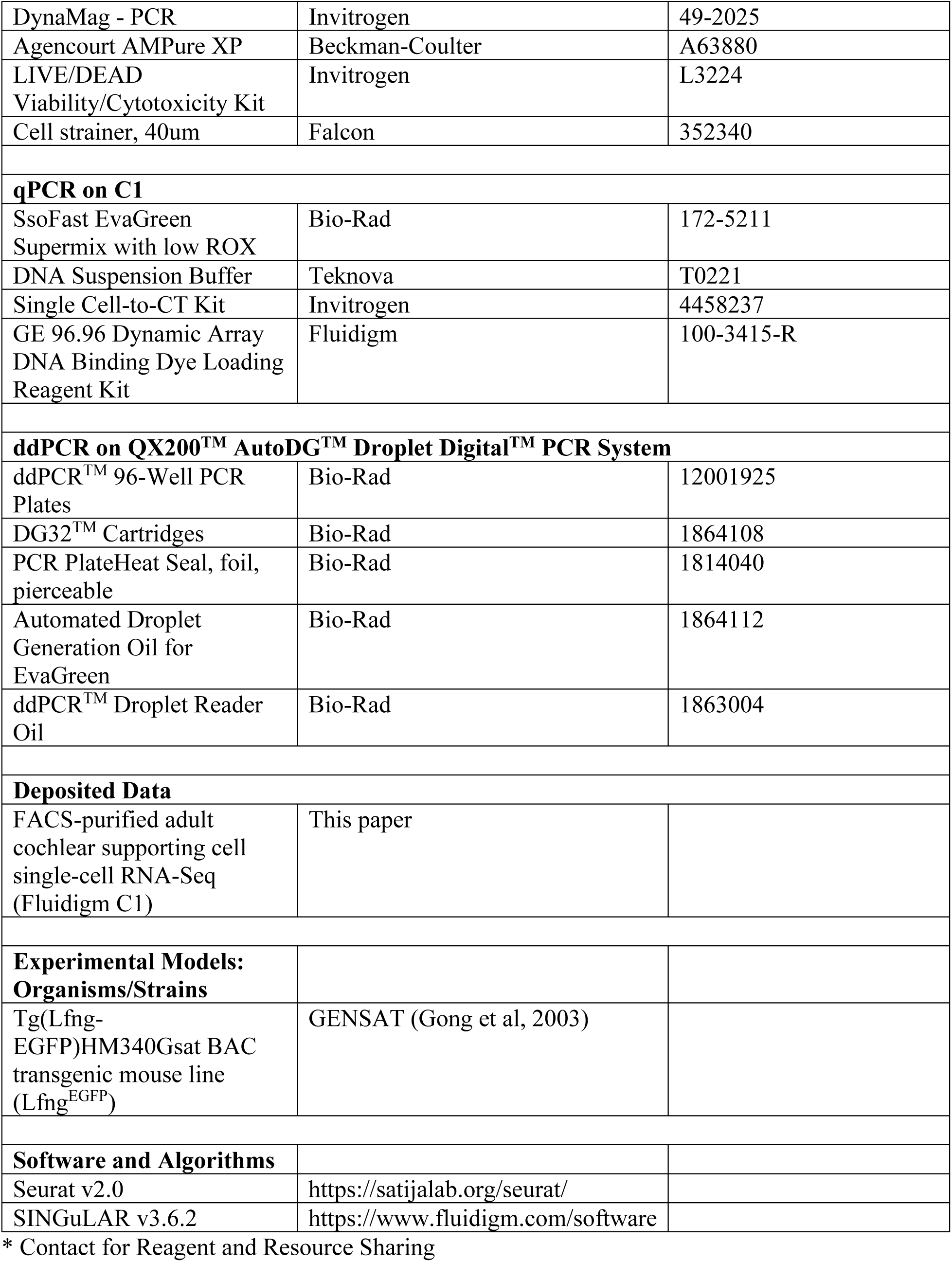
Key Resources.

## Acknowledgments

This research was supported (in part) by the Intramural Research Program of the NIH, NIDCD to M.H. (DC000088), R.J.M. (DC000086), and M.W.K. (DC000059) and NIDCD/NIH Extramural Research Program funds to A.I. (1U24DC015910-01). The authors would like to acknowledge Madeline Pyle, Thomas B. Friedman and Lisa Cunningham who provided helpful feedback and review of this paper. The authors acknowledge Alan Hoofring for his illustrations. We also acknowledge the use of digital droplet PCR resources from T.B.F. (DC000039). This study utilized the high-performance computational capabilities of the Biowulf Linux cluster at the National Institutes of Health, Bethesda, MD. (http://biowulf.nih.gov)

## Ethics Statement

All animal experiments and procedures were performed according to protocols approved by the Animal Care and Use Committee of the National Institute of Neurological Diseases and Stroke and the National Institute on Deafness and Other Communication Disorders, National Institutes of Health.

## Author Contribution Statement

MH and RO contributed to FACS, single cell RNA-sequencing (scRNA-Seq), sc-qPCR and ddPCR. MH, RO, XL, AD, and IT contributed to immunohistochemistry and smFISH. IL, FHL, RO and MH contributed to human temporal bone histopathology. AI analyzed and selected pertinent archival histological material used for immunohistochemical staining and provided support for confocal prescreening of samples. DI and RM were responsible for scRNA-Seq. MH and IT were responsible for scRNA-Seq analysis. MH, DI, RM, MK contributed to writing and revising the manuscript. All authors read and approved final manuscript.

## Supplementary Tables

Supplemental Table S1. Details of single cell captures on Fluidigm C1 capture system.

Supplemental Table S2. Primers utilized for single cell qPCR.

Supplemental Table S3. Primers utilized for ddPCR.

Supplementary Table S4. RNAScope probes.

Supplemental Table S5. Validated Cell Type-Specific Adult Cochlear Supporting Cell Markers.

Supplemental Table S6. Description of validated cell cycle genes expressed by adult cochlear supporting cells.

