## Supplemental Figure S1 for "Characterizing Adult cochlear supporting cell transcriptional diversity using single-cell RNA-Seq: Validation in the adult mouse and translational implications for the adult human cochlea"

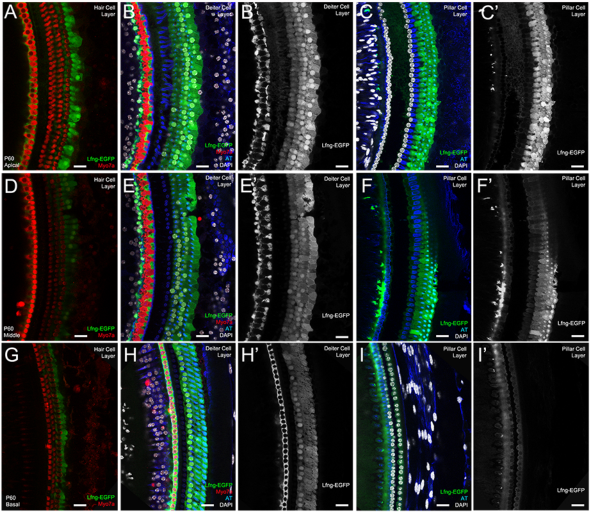


**Supplemental Figure S1. Characterization of Apical to Basal Distribution of Transgene expression in the Adult LfngEGFP cochlea. (A-C'),** Apical turn cochlear whole mounts demonstrate the distribution of GFP expression at the level of the hair cells (A), at the level of Deiters cell layer (B) with single channel GFP grayscale image (B'), and at the level of the pillar cells (C) with single channel GFP grayscale image (C'). **(D-F'),** Middle turn cochlear whole mounts demonstrate the distribution of GFP expression at the level of the hair cells (D), at the level of Deiters cell layer (E) with single channel GFP grayscale image (E'), and at the level of the pillar cells (F) with single channel GFP grayscale image (F'). **(G-I'),** Basal turn cochlear whole mounts demonstrate the distribution of GFP expression at the level of the hair cells (G), at the level of Deiters cell layer (H) with single channel GFP grayscale image (H'), and at the level of the pillar cells (I) with single channel GFP grayscale image (I'). Hair cells are labeled with MYO7A (red), pillar and Deiters cells are labeled with acetylated tubulin (AT, blue), GFP expression (green), and cell nuclei are labeled with DAPI (white). Scale bar, 20 μm.
