## Supplemental Figure S2 for "Characterizing Adult cochlear supporting cell transcriptional diversity using single-cell RNA-Seq: Validation in the adult mouse and translational implications for the adult human cochlea"

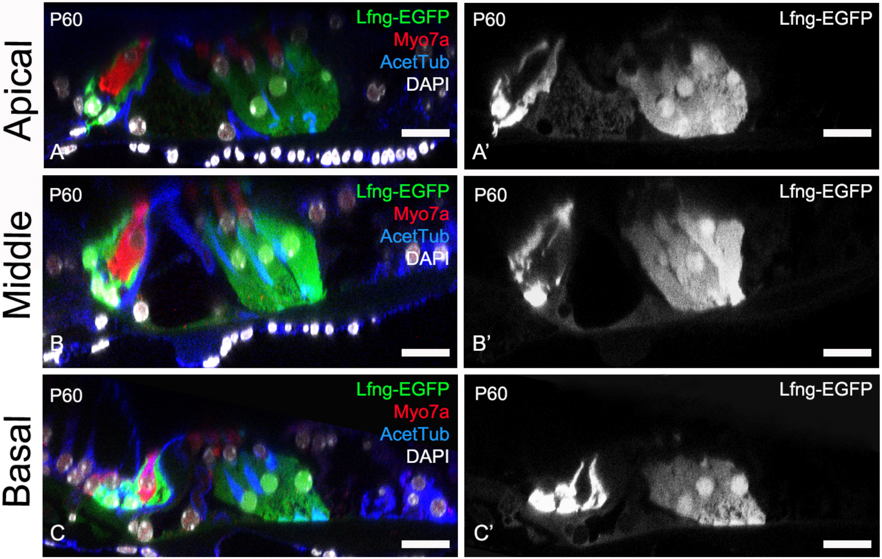


Supplemental Figure S2. Orthogonal reconstructions of Adult LfngEGFP wholemount cochlea. **(A-A'),** Apical turn orthogonal reconstruction of whole mount cochlea (A) with single channel GFP grayscale image (A'). **(B-B'),** Middle turn orthogonal reconstruction of whole mount cochlea (B) with single channel GFP grayscale image (B'). **(C-C'),** Basal turn orthogonal reconstruction of whole mount cochlea (C) with single channel GFP grayscale image (C'). Hair cells are labeled with Myo7a (red), pillar and Deiters cells are labeled with acetylated tubulin (AT, blue), GFP expression (green), and cell nuclei are labeled with DAPI (white). Scale bar, 20 μm.
