## Supplemental Figure S3 for "Characterizing Adult cochlear supporting cell transcriptional diversity using single-cell RNA-Seq: Validation in the adult mouse and translational implications for the adult human cochlea"

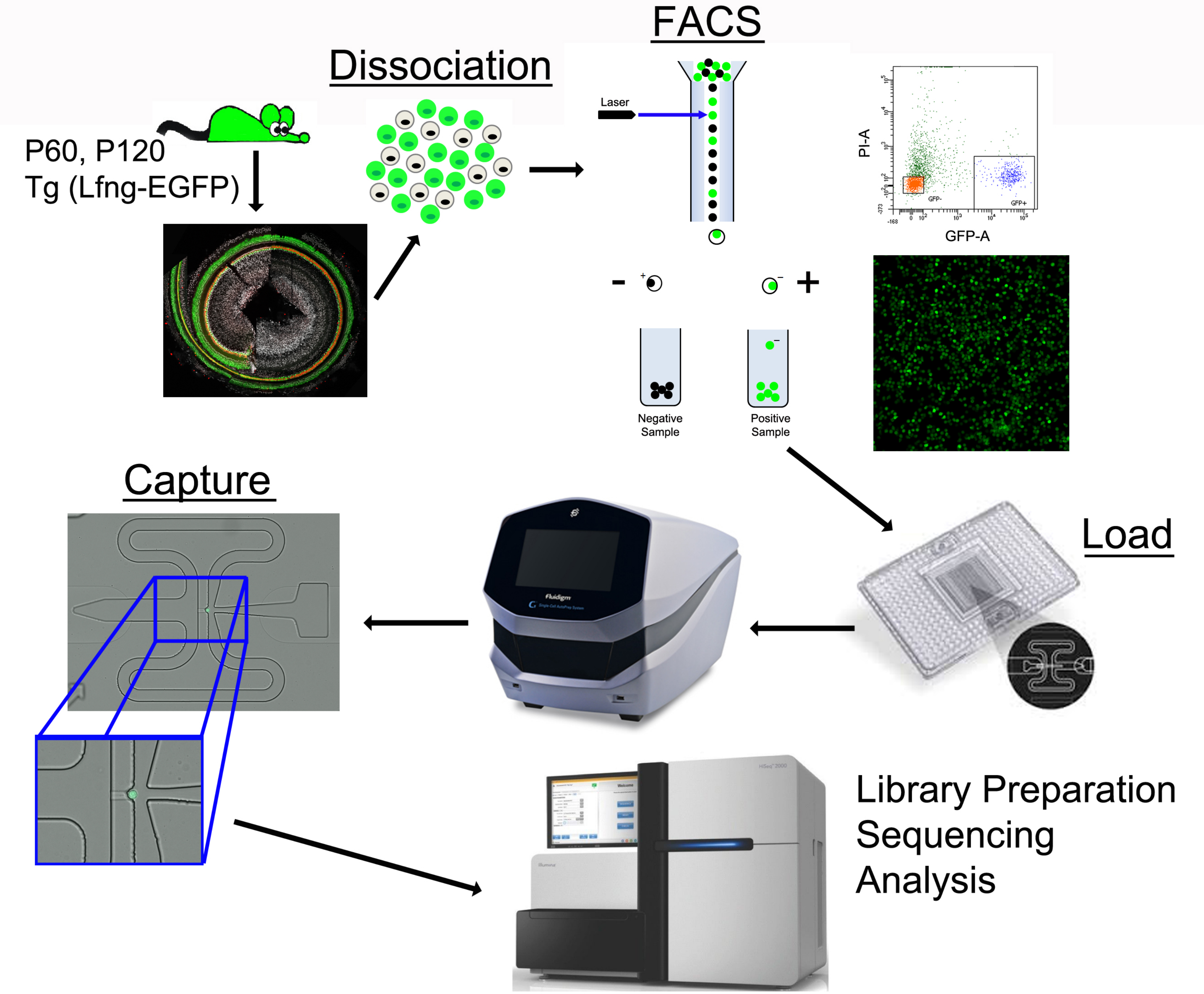


**Supplemental Figure S3. Brief overview of single-cell RNA-Seq workflow of FACS-purified adult cochlear supporting cells utilizing the adult LfngEGFP mouse.** Supporting cells are dissociated and FACS-purified from adult (P60, P120) cochlear supporting cells. FACS-purified LfngEGFP-positive cochlear supporting cells are loaded and captured in individual nests (example of captured LfngEGFP-positive cell shown) in the C1 integrated fluidics chip (IFC), where RNA extraction, reverse transcription and cDNA amplification occur. Libraries were subsequently prepared from the amplified cDNA and sequenced on an Illumina sequencer as noted in the methods.
