## Supplemental Figure S4 for "Characterizing Adult cochlear supporting cell transcriptional diversity using single-cell RNA-Seq: Validation in the adult mouse and translational implications for the adult human cochlea"

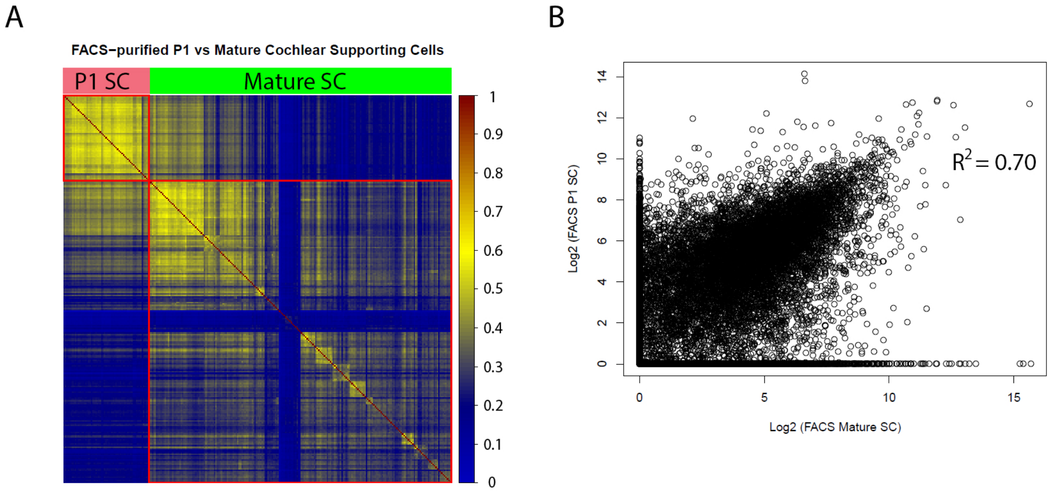


**Supplemental Figure S4. Correlation analysis between P1 and Mature (P60, P120) LfngEGFP-positive cochlear supporting cells. (A),** The non-averaged correlogram highlights the distinct gene expression profile of mature cochlear supporting cells. Cells are arrayed along the x- and y-axis in the same order and the individual squares represent the Spearman correlation between the gene expression in between each pair of cells. Note the large cell-to-cell variability in gene expression between P1 cochlear supporting cells (red bar) and mature cochlear supporting cells (green bar). Scale for degree of correlation ranges between 0 (no correlation, blue) to complete correlation (complete correlation, dark orange). Red boxes highlight the greater degree of dissimilarity between P1 and mature SCs by demonstrating the higher degree of correlation among mature SCs and P1 SCs, respectively. **(B),** Correlation plots comparing the average gene expression across all mature cochlear supporting cells (x-axis) to the average gene expression across all P1 cochlear supporting cells (along the y-axis). Expression values along both axes are in Log2 [TPM].
