## Supplemental Figure S5 for "Characterizing Adult cochlear supporting cell transcriptional diversity using single-cell RNA-Seq: Validation in the adult mouse and translational implications for the adult human cochlea"

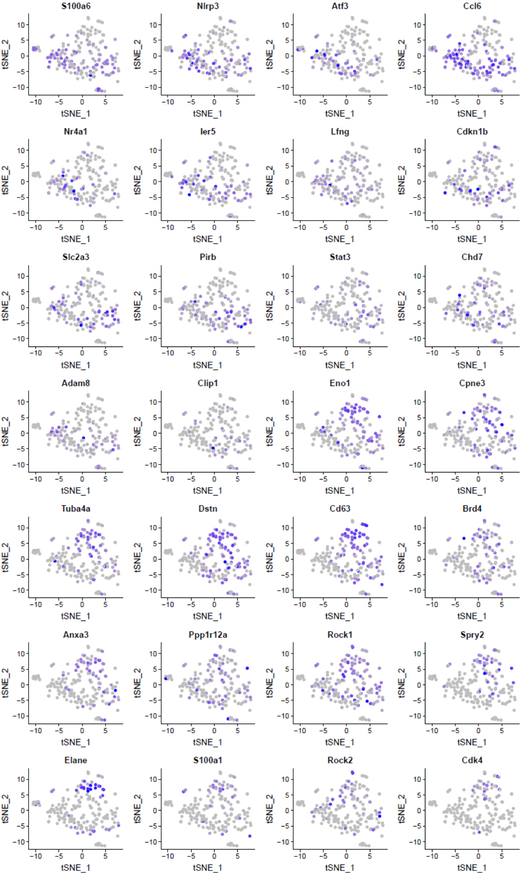


**Supplemental Figure S5.** Adult cochlear SC candidate genes identified from scRNA-Seq. Feature plots demonstrate distribution of expression across FACS-purified adult cochlear SCs obtained from the Fluidigm C1 platform.
