## Supplemental Figure S6 for "Characterizing Adult cochlear supporting cell transcriptional diversity using single-cell RNA-Seq: Validation in the adult mouse and translational implications for the adult human cochlea"

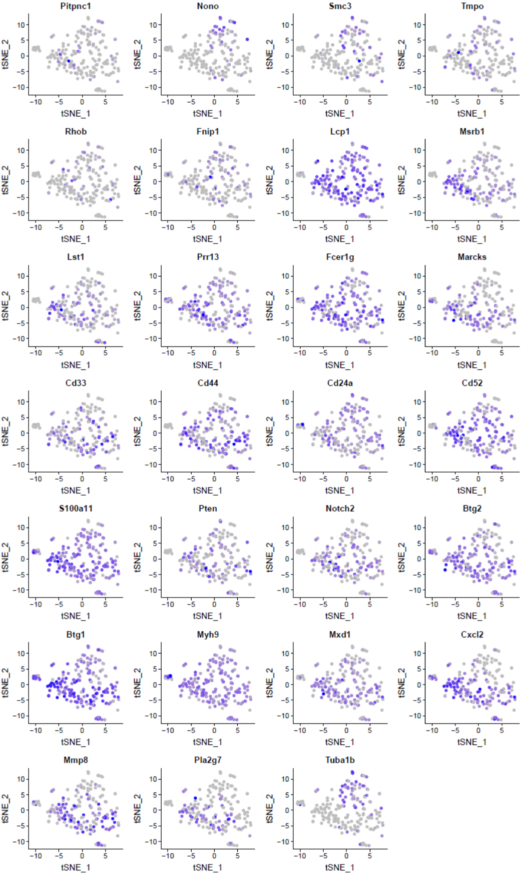


**Supplemental Figure S6.** Adult cochlear SC candidate genes identified from scRNA-Seq (cont'd from Supplemental Figure S5). Feature plots demonstrate distribution of expression across FACS-purified adult cochlear SCs obtained from the Fluidigm C1 platform.
