## Supplemental Figure S7 for "Characterizing Adult cochlear supporting cell transcriptional diversity using single-cell RNA-Seq: Validation in the adult mouse and translational implications for the adult human cochlea"

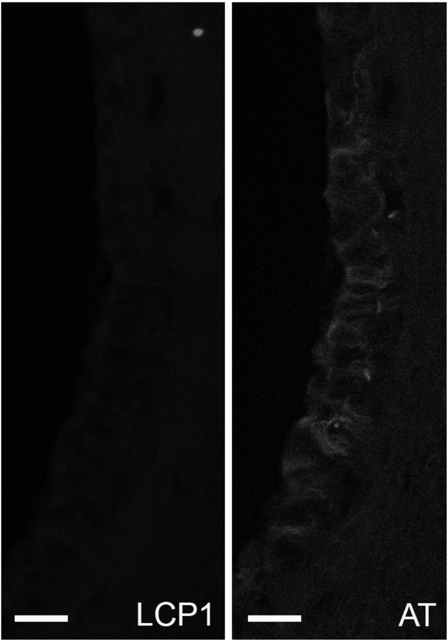


**Supplemental Figure S7.** Expression of Acetylated Tubulin and LCP1 in the human stria vascularis. Unlike acetylated tubulin (AT), LCP1 is not expressed in the stria vascularis in the human inner ear. Grayscale single channel images shown for LCP1 (Right) and AT (Left). Scale bar, 20 μm.
