## Supplemental Figure S9 for "Characterizing Adult cochlear supporting cell transcriptional diversity using single-cell RNA-Seq: Validation in the adult mouse and translational implications for the adult human cochlea"

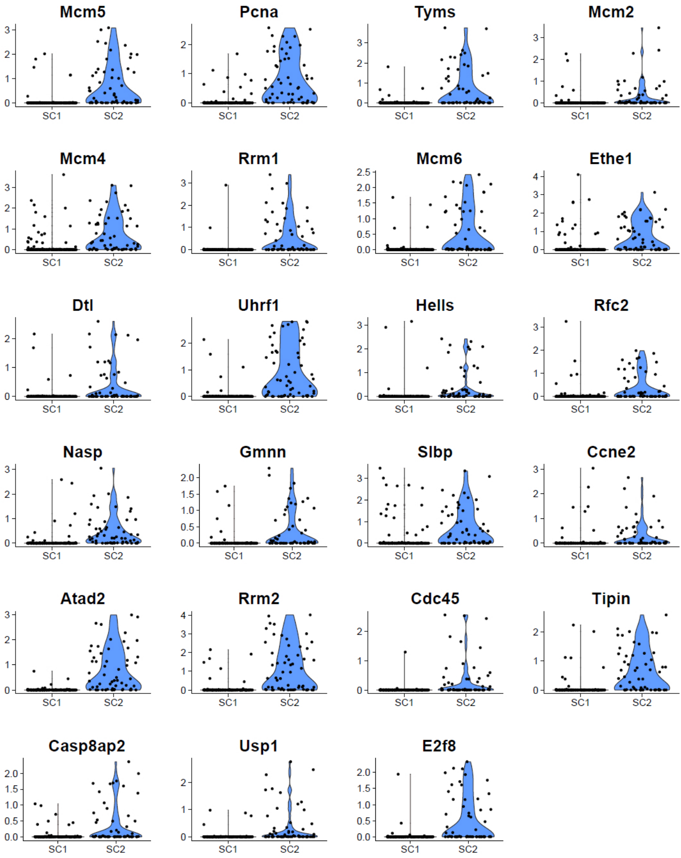


**Supplemental Figure S9. Cell cyle: S phase-associated gene expression as delineated by Nestorowa and colleagues in FACS-purified adult cochlear supporting cells.** Violin plots of scRNA-Seq data with supporting cell clusters on the horizontal axis and gene expression level in log2[TPM] demonstrate expression in adult cochlear supporting cells.
