## Supplemental Figure S10 for "Characterizing Adult cochlear supporting cell transcriptional diversity using single-cell RNA-Seq: Validation in the adult mouse and translational implications for the adult human cochlea"

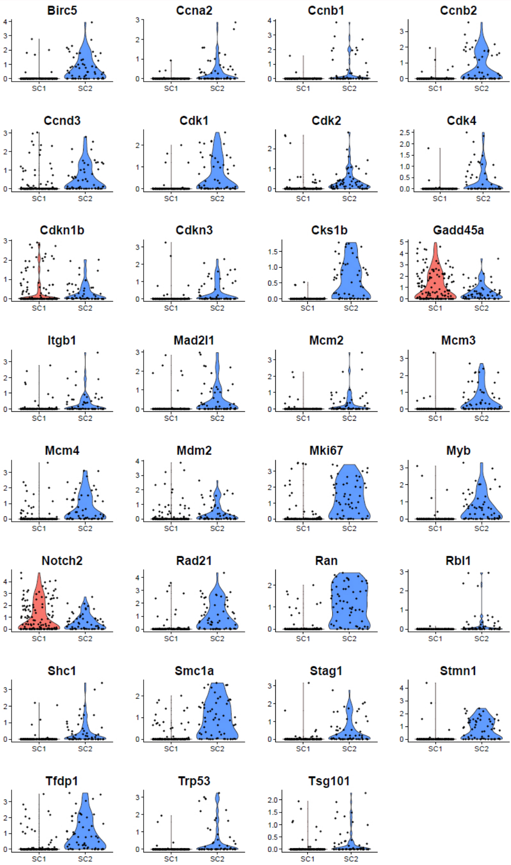


Supplemental Figure S10. Subset of cell cycle gene expression retained by Lgr5-positive neonatal SCs as delineated by Cheng and colleagues that are expressed by FACS-purified adult cochlear SCs. Violin plots of scRNA-Seq data with SC clusters on the horizontal axis and gene expression level in log2[TPM] demonstrates expression in adult cochlear SCs. The SC2 cluster of adult cochlear supporting cells appears to express a large proportion of these genes. Genes including Cdkn1b, Gadd45a and Notch2 are expressed by both supporting cell clusters.
