## Supplemental Table S1 for "Characterizing Adult cochlear supporting cell transcriptional diversity using single-cell RNA-Seq: Validation in the adult mouse and translational implications for the adult human cochlea"

**Supplemental Table S1. Details of single cell captures on Fluidigm C1 capture system.**

| **Isolation #** | **1 (9c)** | **2 (10c)** | **3 (17c)** | **4 (19c)** | **5 (20c)** | **6 (21c)** | **7 (28)** | **8 (31)** | **9 (32)** | **10 (36)** | **11 (38)** | **12 (40)** |
| --- | --- | --- | --- | --- | --- | --- | --- | --- | --- | --- | --- | --- |
| **Organ** | **P60**  **Cochlea** | **P60**  **Cochlea** | **P60**  **Cochlea** | **P60**  **Cochlea** | **P60**  **Cochlea** | **P60**  **Cochlea** | **P120**  **Cochlea** | **P60**  **Cochlea** | **P120**  **Cochlea** | **P120**  **Cochlea** | **P120**  **Cochlea** | **P60**  **Cochlea** |
| **Method** | **FACS** | **FACS** | **FACS** | **FACS** | **FACS** | **FACS** | **FACS** | **FACS** | **FACS** | **FACS** | **FACS** | **FACS** |
| **# Mice** | **4** | **4** | **4** | **4** | **4** | **4** | **4** | **4** | **4** | **4** | **4** | **4** |
| **Genotype** | **Lfng^EGFP^** | **Lfng^EGFP^** | **Lfng^EGFP^** | **Lfng^EGFP^** | **Lfng^EGFP^** | **Lfng^EGFP^** | **Lfng^EGFP^** | **Lfng^EGFP^** | **Lfng^EGFP^** | **Lfng^EGFP^** | **Lfng^EGFP^** | **Lfng^EGFP^** |
| **# single-cell captures** | **37** | **24** | **4** | **13** | **13** | **12** | **47** | **40** | **33** | **18** | **34** | **33** |
| **# multicell captures** | **3** | **0** | **0** | **0** | **0** | **0** | **0** | **0** | **0** | **0** | **0** | **0** |
| **# GFP** | **35** | **23** | **4** | **13** | **13** | **12** | **47** | **40** | **33** | **18** | **34** | **33** |
| **# negative** | **2** | **1** | **0** | **0** | **0** | **0** | **0** | **0** | **0** | **0** | **0** | **0** |
| **# single cells sequenced** | **33** | **21** | **3** | **13** | **13** | **11** | **47** | **27** | **29** | **17** | **32** | **33** |
| **# outliers** | **14** | **8** | **0** | **6** | **5** | **6** | **10** | **4** | **2** | **1** | **3** | **9** |
| **# single cells analyzed** | **19** | **13** | **3** | **7** | **8** | **5** | **37** | **23** | **27** | **16** | **29** | **24** |
| **Sequencing lanes** | **1** | **1** | **1** | **1** | **1** | **1** | **2,3** | **2,3** | **2,3** | **2** | **2** | **2** |
