## Supplemental Table S2 for "Characterizing Adult cochlear supporting cell transcriptional diversity using single-cell RNA-Seq: Validation in the adult mouse and translational implications for the adult human cochlea"

**Supplemental Table S2. Primers utilized for single cell qPCR.**

| Gene | Forward primer (5’-3’) | Reverse primer (5’-3’) |
| --- | --- | --- |
| *Adam8* | AGGTGCTGTGTCCAGTTCA | TGGTGGGAGGATCAGGTCTA |
| *Alyref* | TGCCCTGAAGGCTATGAAACA | TGCAGGTCTTCGCTGTGTA |
| *Anxa3* | TCTTGTGTCTACGGAGCTTCC | GTCCTCAATGTCCTTCTGGCTA |
| *Atf1* | TGCATCTCAGCCTGGTTCAA | ATGAGTCCTGAGACTCCTCAC |
| *Atf2* | AGCCACCTCCACTACAGAAAC | TTCTTCGACGGCCACTTGTA |
| *Atf3* | GCTGCCAAGTGTCGAAACAA | AGCTCAGCATTCACACTCTCC |
| *Birc5* | AGGAATTGGAAGGCTGGGAA | GCCAGGGGAGTGCTTTCTA |
| *Braf* | GGAGAGCATAACCCACCATCA | GCTGCTGTTCTCTTTGCTGAA |
| *Brd4* | CCCCTGACCATGAAGTGGTA | CTCATCAGGCATCTTGGCAAA |
| *Cbl* | ACCCAGAGCAGATGTCCCTA | CTGCTACTGGGCTGCTATTCA |
| *Ccl6* | ATCGTCGCTATAACCCTCCA | CATGGGATCTGTGTGGCATAA |
| *Ccnb2* | CATGGCCAAGAACGTTGTGAA | GGAGTCTGCTGCTGGCATA |
| *Ccnd3* | ATGCGGAAGATGCTGGCATA | GTTCATAGCCAGAGGGAAGACA |
| *Cd63* | GGGCCTGCAAGGAGAACTAC | GCCACCTCCACAAGCATGATA |
| *Cdk1* | TGCCAGAGCGTTTGGAATAC | CACTTCTGGAGATCGGTACCA |
| *Cdk2* | GGAGAAGTTGTGGCGCTTAA | GAGATCTCTCGGATGGCAGTA |
| *Cdk4* | GCTGACTTTGGCCTAGCTAGAA | ACTTCAGGAGCTCGGTACCA |
| *Cdkn1b* | CAGTGTCCAGGGATGAGGAA | TTCGGGGAACCGTCTGAAA |
| *Cdkn3* | CAGACGAAGAACCTGTTGATGAA | ATTCACTCGCGACAGAGGTA |
| *Cenpx* | AGCTCATGGCGGAGTTCC | TGATCCACTTCCACAACATCCA |
| *Cflar* | TGCCTGAAGAACATCCACAGAA | GCGAAGCCTGGAGAGTATTCATA |
| *Chd7* | CCAACGAGAGCACCATTCAA | CACCTCCATGGACTCTTCCTTA |
| *Cks1b* | AGTCAGGGATGGGTCCACTA | GTGGCCGTCGGAACAGTA |
| *Clec7a* | ACTTCAGCACTCAAGACATCCA | AATGGGCCTCCAAGGTGAA |
| *Cpne3* | TGACTGCCATCTGGTCTGTA | CCTGAGCTCCAAAGCCAAAA |
| *Csnk1a1* | GGGTATTGGGCGTCACTGTAA | GCCTTGTCCTGTTGTCTCTGTA |
| *Csnk2a1* | CTCAGCACTTCCTGTGTCTGTA | CTGCCAACACAGCTGTTTCA |
| *Ddost* | TACGGGGAGTTCCTCTATGACA | CACGTTGATGTTGCCTCCAAA |
| *Dstn* | ACCAGAACAAGCACCTCTGAA | GGCCCATTTGCTTGATACTCA |
| *Dusp6* | GCTGCTGCTCAAGAAACTCA | TCGGCCTGGAACTTACTGAA |
| *Eef1g* | CTGTCGTGCGTATGCCTTTTA | AGGAGACAAGGGAGGAGGAA |
| *Eif4e* | TAAGGCACAGTTGCTCAGTCC | GATGCACCAAAACCCCAATCC |
| *Elane* | ATGGCTTTGACCCATCACAAC | GCACGTTGGCGTTAATGGTA |
| *Eno1* | CGCCTGGCCAAGTACAATCA | CTGAAGGACCTGCCAGCAAA |
| *Ep300* | CAAACATGCCTTTGGCTCCA | AGAGTTCACTGGGCAAGAAGAA |
| *Fnip1* | GTGTGGGCATGTTGGCAAA | AGGAGATCCAATAGCGCCATAC |
| *Fos* | ATGGGCTCTCCTGTCAACAC | GCTGTCACCGTGGGGATAAA |
| *Gadd45a* | CCTGCACTGTGTGCTGGT | TGATCCATGTAGCGACTTTCCC |
| *Grb2* | TCAACATCCGTGTCCAGGAA | AAAGAGCGCCTGGACGTA |
| *Gsk3b* | GCAGCCTTCAGCTTTTGGTA | GGAGTTGCCACTACTGTGGTTA |
| *Ier5* | GAGGAGATGGAGACCGGGAA | TTTCCGTAGGAGTCCCGAGAA |
| *Jun* | GGAACAGGTGGCACAGCTTA | CGTTTGCAACTGCTGCGTTA |
| *Kif15* | ACAGCTGAGTGACCTGGAAA | GGTCATGCACCTCACATTTCA |
| *Kif20b* | AGTATTGGAGCCAACGGGAA | CGACGTTCAGCGTTTTCTTCTA |
| *Lcp1* | GGATCCGTGTCTGACGAAGAA | GCTCATTGCAGCTGATGTATCC |
| *Lfng* | TCGATCTGCTGTTCGAGACC | CCTCCCCATCAGTGAAGATGAA |
| *Lsm3* | AGCCTGGATGAGCGAATTTA | TCTTCTACATCTCCCAGGATCA |
| *Mapk1* | CGTTGGTACAGAGCTCCAGAA | TGCAGCCCACAGACCAAATA |
| *Mapk3* | TGGGCCAAGCTCTTTCCTAA | TGCGCTTGTTTGGGTTGAA |
| *Mcm3* | ACCGATGATTCTCAGGAGACC | GCCGCCTTAAAAGCCTTCA |
| *Mcm4* | TCTGCAACTGACCCTCGTA | GTTTACGAGAAGTGGCACTCA |
| *Mdm2* | GCGCAAAACGACACTTACACTA | TGCTGCTGCTTCTCGTCATA |
| *Mki67* | GAGACATACCTGAGCCCATCA | GCTTTGCTGCATTCCGAGTA |
| *Mrpl18* | TAGGGGTAGCTCGGAAAGAAC | GGCCGTTAAGATGCTCAACA |
| *Myb* | TCCTCCGTCAACAGCGAATA | CAATGCGACAGGATAGGGAAC |
| *Nfkb1* | ACCGTATGAGCCTGTGTTCA | GTAGCCTCGTGTCTTCTGTCA |
| *Nlrp3* | TGCTCTGCAACCTCCAGAAA | AACCAATGCGAGATCCTGACA |
| *Nono* | ACGTCATGAGCACCAGGTTA | TATGCAGCTCCTCCATTCTCC |
| *Nras* | AGCAGGTGGTGTTGGGAAAA | GGCAGGTCTCACCATCAATCA |
| *Park7* | TGGATGCAAGGTCACAACAC | CGGCTCTCTGAGTAGCTGTA |
| *Pirb* | GAGGATGGAGTGGAGCTGAA | AGGGTTTCACCTGGGCATAA |
| *Pitpnc1* | CTGCCGAAATTCTCCATCCA | GGTCTTTGGCTTCACTGTCA |
| *Ppib* | TCCGTGGCCAACGATAAGAA | CTCGTCCTACAGATTCATCTCCAA |
| *Psenen* | TCGTGATCTTGCGTCTGTCA | GGATACCCGCTCCAAGTTCATA |
| *Psmb1* | GGTTTTCGCCTTATGCCTTCA | TGTCTGAAGCGACGATGGAA |
| *Psmb2* | GAGGGCAGTGGAGCTTCTTA | ACTGAAGGTGGGCAGATTCAA |
| *Ptgs2* | CTTCTCCCTGAAGCCGTACA | TGTCACTGTAGAGGGCTTTCAA |
| *Rad21* | GCACTCAGCAGATGCTTCA | TCGACACAGCTCAAGCAAAC |
| *Ran* | TGCGCGATGGCTACTACA | GGATGTTTTCACACACTCGTACC |
| *Rock1* | GCAGAGGTGCATTTGGAGAA | GCTGAGCAGCTTCATAGCATA |
| *Rock2* | GTGGGAAATCAGCTGCCTTTTA | TTCTCTACAAGGTGGAGAGTCAC |
| *Rpl10a* | CTGCCTTCTTTCCGGTTTCC | TTCCCGTGCAGGACTTCC |
| *Rpl36a* | TAAACTTCCGTTGCGGCTCTA | AGAATGTCCGGCGGGTTTTA |
| *Rpl36al* | GCTCTGTTCGTCCCTTTCC | ACGTTGACCATGTTTGCAGTA |
| *Rpl39* | GGCCTTTCTCTTCTCCATTCC | CATCCGGATCCACTGAGGAA |
| *Rps20* | GTTCGCTCCTGCTGAGGAA | GGGCGTCTTTCCGGTATCTTTA |
| *S100a1* | TGCCATGGAGACCCTCATCA | TTGTCCACAGCATCTGCATCC |
| *S100a6* | AAGCACACCCTGAGCAAGAA | AGCCTTGCAATTTCAGCATCC |
| *Shc1* | TTGCCAACCATCACATGCA | GGCAACATAGGCAACATACTCA |
| *Slc2a3* | TCTGGTCGGAATGCTCTTCC | GAGGAAGGCAGCGAAGATGATA |
| *Smc1a* | TTGACGCTGCCTTGGATAACA | GCCTGGAAGTTGCAAGTTGAC |
| *Smc3* | GGCAGAAATGGCTCTGGAAA | TCTGGACGAAGATGGCTGAA |
| *Snrpd2* | TGTCGCAACAACAAGAAGCT | CCTCAGTCCACATCTCCTTCA |
| *Spcs1* | TCGGTATCGGTCTTCAGCAA | AGCTAGTTTCTGCCCCTTGTA |
| *Spry2* | GGAGAGGGGTTGGTGCAAA | AGGTCTTGGCAGTGTGTTCA |
| *Srp19* | AGGGGAGACGGATCCCTATAA | CAGTCCAACTGCTGAGCATAC |
| *Srsf2* | CAAGAGCCCACCCAAGTCT | GTTAAGCCGCTTGCCGATT |
| *Ssr2* | TCTCCGGTATGCTCAATGTCAA | TTTGAGAGGACGCAGGACAA |
| *Ssr4* | GCCGACGTTAGTGGAAAACA | TTGTGCTCCAGGCTCCA |
| *Stag1* | TTGACTGGTTTGTCAGACTCC | CAGAGCAGTCATCAGCTTCA |
| *Stat3* | TGGGCATCAATCCTGTGGTA | CCAATTGGCGGCTTAGTGAA |
| *Stmn1* | TCTGTTGGTGCTCAGAGTGT | CTACACAATCCACTGGCAAGG |
| *Tbrg1* | CTCCTCTGCAGATGCCTGTTA | TGAAAGTGGGTTGGGCATCA |
| *Tfdp1* | AGGTGGCCAGAAGTTTGGTA | TCGTGACGAATACTCCACCAA |
| *Tmpo* | GGAGTGAATCCTGGTCCCATT | TCCCTCAGCTTCAACAGCTT |
| *Tuba4a* | CTTTGTGGACCTGGAGCCTA | ATAAGCTGCTCTGGGTGGAA |
