## Supplemental Table S3 for "Characterizing Adult cochlear supporting cell transcriptional diversity using single-cell RNA-Seq: Validation in the adult mouse and translational implications for the adult human cochlea"

**Supplemental Table S3. Primers utilized for ddPCR.**

| Gene | Forward primer (5’-3’) | Reverse primer (5’-3’) |
| --- | --- | --- |
| *Atf3* | TGCATCTCAGCCTGGTTCAA | ATGAGTCCTGAGACTCCTCAC |
| *Btg1* | GGGATCAGGTTACCGTTGTATT | CCCAAGAGAAGTTCCTCCTTAC |
| *Btg2* | CTGCTTTGTATGGGTGGATAGT | GAGCCACCTCTCAAAGGAATAG |
| *Ccnd3* | ATGCGGAAGATGCTGGCATA | GTTCATAGCCAGAGGGAAGACA |
| *Cdkn1b* | CAGTGTCCAGGGATGAGGAA | TTCGGGGAACCGTCTGAAA |
| *CD44* | CAGTCACAGACCTACCCAATTC | GTGTGTTCTATACTCGCCCTTC |
| *CD52* | TTCCTCCTCTTCCTCACTATCA | TTGGTGGAGGTGCTGTTT |
| *Fcer1g* | CCTGGATGCTGTCCTGTTT | AGAGGGCTCGGAGAGAATTA |
| *Lcp1* | GGATCCGTGTCTGACGAAGAA | GCTCATTGCAGCTGATGTATCC |
| *Lfng* | TCGATCTGCTGTTCGAGACC | CCTCCCCATCAGTGAAGATGAA |
| *Msrb1* | CAAACTCATCTTGCCTCACTCTA | CTGGACAAAGTGGTGAGAAGAG |
| *Mxd1* | CCATACAGCAAGGACAGAGATG | CCGCTGAAGCTGGTCTATTT |
| *Myh9* | TGCCCTGGAACTGTGTTTAG | GTCCCACTTGCTCCTTTATGA |
| *Nlrp3* | TGCTCTGCAACCTCCAGAAA | AACCAATGCGAGATCCTGACA |
| *Notch2* | AGACTGGCGACTTCACTTTC | TCCACACAAACTCCTCCATTC |
| *Prrl13* | GAATAGGAGTGACCAGGAAGTG | TATGGATTCGGTCCAGCATTAG |
| *Pten* | CCCACCACAGCTAGAACTTATC | GGGTCTGTAATCCAGGTGATTT |
| *S100a1* | TGCCATGGAGACCCTCATCA | TTGTCCACAGCATCTGCATCC |
| *S100a6* | AAGCACACCCTGAGCAAGAA | AGCCTTGCAATTTCAGCATCC |
| *S100a11* | GAGATGCATTGAGTCCCTGATT | TAAGCCACCAATGAGGTTGAG |
| *Slc2a3* | TCTGGTCGGAATGCTCTTCC | GAGGAAGGCAGCGAAGATGATA |
| *Spry2* | GGAGAGGGGTTGGTGCAAA | AGGTCTTGGCAGTGTGTTCA |
