## Supplemental Table S4 for "Characterizing Adult cochlear supporting cell transcriptional diversity using single-cell RNA-Seq: Validation in the adult mouse and translational implications for the adult human cochlea"

**Supplementary Table S4. RNAScope probes**

| Gene | Accession Number | Probed Region (nt) | ZZ Oligo pairs |
| --- | --- | --- | --- |
| *S100a6* | NM_011313.2 | 8-674 | 13 |
| *Lcp1* | NM_001247984.1 | 1230-2350 | 20 |
| *Nlrp3* | NM_145827.3 | 2 – 1098 | 20 |
| *Slc2a3* | NM_011401.4 | 503 – 1665 | 20 |
| *Spry2* | NM_011897 | 2 – 905 | 20 |
| *Birc5* | NM_001012273.1 | 131 – 1036 | 20 |
| *Notch2* | NM_010928.2 | 215 – 1068 | 20 |
| *Tuba1b* | NM_006082.2 | 107-1729 | 6 |
| *Myh9* | NM_022410.3 | 240-3331 | 20 |
| *Cdkn1b* | NM_009875.4 | 339-1708 | 20 |
| *Pla2g7* | NM_013737.5 | 670-1703 | 20 |
| *Ppib* | NM_011149.2 | 98-856 | 15 |
| *Dap8* | EF191515 | 414-862 | 10 |
