## Supplemental Table S5 and S6 for "Characterizing Adult cochlear supporting cell transcriptional diversity using single-cell RNA-Seq: Validation in the adult mouse and translational implications for the adult human cochlea"

**Supplemental Table S5: Validated Cell Type-Specific Adult Cochlear Supporting Cell Markers**

| Gene Symbol | Gene Name | Organ of Corti Localization  (Adult Mouse) | Possible functional roles | References demonstrating presence in the inner ear |
| --- | --- | --- | --- | --- |
| Lcp1 | Lymphocyte cytosolic protein 1 | Pillar, Deiters cells | Actin-bundling protein (Daudet et al, 2002; Todd et al, 2011) | This publication |
| S100a6 | Calcyclin | Medial supporting cells, Pillar cells and Deiters cells | Calcium binding protein implicated in cell proliferation, cytoskeletal dynamics, and tumorigenesis (Lesniak et al, 2009) | This publication |
| Myh9 | Myosin heavy chain 9 | Adult organ of Corti hair cells and supporting cells | Cytokinesis, cell motility, and maintenance of cell shape; implicated in localization to the nucleus, antiproliferative activity, tumor suppressor activity (Cho et al, 2012; Coaxum et al, 2017) | Mhatre et al, 2004 |
| Nlrp3 | Cryopyrin | Medial supporting cells, pillar cells, Deiters cells | Regulation of cochlear autoinflammation (Nakanishi et al, 2017) | This publication |
| Slc2a3 | Solute carrier 2 member 3 or GLUT3 | Pillar cells | Glucose transport (Trattner et al, 2013; Patching, 2017) | This publication |
| Cdkn1b | Cyclin-dependent kinase inhibitor 1b | Pillar cells, Lateral supporting cells | Cell cycle control | Laine et al, 2010;  Oesterle et al, 2011 |
| Tuba1b | Acetylated tubulin | Pillar cells, Lateral supporting cells | Microtubule component, Cytoskeletal dynamics | Lowenheim et al, 1999; Oesterle et al, 2009 |
| Spry2 | Sprouty2 | Deiters cells | Role in establishing normal pillar cell cytoarchitecture by antagonizing FGF signaling during early postnatal cochlear development (Shim et al, 2008) | Shim et al, 2008 |
| Pla2g7 | Phospholipase A, Group VII | Medial supporting cells | Role unknown | This publication |
| S100a11 | S100a11 calcium binding protein | Medial supporting cells, Pillar cells, some Deiters cells | Double-stranded DNA repair, cell cycle regulation & proliferation, role in inflammation (Foertsch et al, 2016; Huang et al, 2016) | This publication |
| S100a1 | S100a1 calcium binding protein | Medial supporting cells, Deiters cells | Calcium-binding protein implicated in cytoskeletal dynamics in the adult organ of Corti (Coppens et al, 2001) | Coppens et al, 2001 |

**Supplemental Table S6: Validated cell cycle genes expressed by adult cochlear supporting cells**

| Gene Symbol | Gene Name | Gene Ontology: Biological Process | Possible functional roles | Reference |
| --- | --- | --- | --- | --- |
| Birc5 | Survivin | positive regulation of transcription from RNA polymerase II promoter | Possible promotion of cell proliferation, prevention of apoptosis | Froehlich et al, 2015; Habtemichael et al, 2010 |
| Ccna2 | Cyclin A2 | positive regulation of transcription from RNA polymerase II promoter | Promotes transition through B1/S and G2/M phase | Chen et al, 2009; Zhong et al, 2015 |
| Ccnb1  Ccnb2 | Cyclin B1  Cyclin B2 | positive regulation of transcription from RNA polymerase II promoter | Form maturation-promoting factor (MPF); cell cycle regulation | Lu et al, 2013 |
| Ccnd3 | Cyclin D3 | positive regulation of transcription from RNA polymerase II promoter | Regulates transition through G1/S phase of cell cycle | Sherr, 1995; Lee et al, 2017; Laine et al, 2010 |
| Cdkn3 | Cyclin-dependent kinase inhibitor | positive regulation of transcription from RNA polymerase II promoter | Upregulation associated with downregulation of Cdkn1b, resulting in increased cell proliferation | Wang et al, 2017 |
| Cks1b | Cdc28 protein kinase regulatory subunit 1b | positive regulation of transcription from RNA polymerase II promoter | E3 ligase that ubiquitylates substrates, designating them for proteosomal degradation | Pavlides et al, 2016 |
| Itgb1 | Beta1-integrin | positive regulation of transcription from RNA polymerase II promoter; negative regulation of DNA endoreduplication | Critical to laminin-sonic hedgehog-induced cell proliferation | Blaess et al, 2004 |
| Myb | Myb proto-oncogene | positive regulation of transcription from RNA polymerase II promoter | Regulation of epithelial to mesenchymal transition | Kurima et al, 2011; Tanno et al, 2010 |
| Notch2 | Notch 2 | positive regulation of transcription from RNA polymerase II promoter | Regulation of cell-fate determination | Ohashi et al, 2017; Teng et al, 2017 |
| Ran | Ran encodes Ras-related nuclear protein | positive regulation of transcription from RNA polymerase II promoter | Small GTPase involved in nucleocytoplasmic transport, DNA synthesis and cell cycle progression | Deng et al, 2013 |
| Rbl1 | RB transcriptional co-repressor like-1 | positive regulation of transcription from RNA polymerase II promoter | Cell cycle regulator, regulates G1/S phase transition | Yu et al, 2010 |
| Stmn1 | Stathmin 1 | positive regulation of transcription from RNA polymerase II promoter | Ubiquitous cytosolic phosphoprotein which prevents microtubule assembly (Rubin & Atweh, 2004); Associated with increased proliferation and poor prognosis in variety of human malignancies and implicated as marker of PI3K pathway activation | Jiang et al, 2018 |
| Tfdp1 | Transcription factor Dp-1 | positive regulation of transcription from RNA polymerase II promoter | Enhances DNA binding and transcription of E2F target genes | Hitchens & Robbins, 2003 |
| Tsg101 | Tumor susceptibility gene 101 | positive regulation of transcription from RNA polymerase II promoter | Suppress p21^CIP1/WAF1^, a cyclin-dependent kinase inhibitor which negative regulates the cell cycle | Lin et al, 2013 |
| Cdk1  Cdk4  Cdk6 | Cyclin-dependent kinases 1, 4, and 6 | G1/S transition of mitotic cell cycle | Kinases essential for cell cycle G1 phase progression | Malgrange et al, 2003 |
| Gadd45a | Growth arrest and DNA damage alpha | negative regulation of DNA endoreduplication | Facilitates somatic cell reprogramming by relaxing chromatin structure around the miR-295 promoter | Hu et al, 2009; Li et al, 2017 |
| Mad2l1 | Mitotic arrest deficient 2 like 1 | negative regulation of DNA endoreduplication | Mitotic delay; prevents anaphase progression until all kinetochores are bound by microtubules | Marks et al, 2017 |
| Mcm2 | Minichromosome maintenance complex 2 | negative regulation of DNA endoreduplication | Component of minichromosome maintenance complex (MCM), a DNA helicase critical for initiating DNA replication; missense variant implicated in autosomal dominant nonsyndromic deafness | Zhai et al, 2017; Gao et al, 2015 |
| Mdm2 | Mouse double minute 2 | negative regulation of DNA endoreduplication | E3 ubiquitin ligase that ubiquitinates downstream targets including p53 and Rb, which then targets them for proteosomal degradation; promotes genomic instability and tumorigenesis by delaying p53-independent DNA breakage repair | Saadatzadeh et al, 2017; Laos et al, 2017 |
| Mki67 | Marker of proliferation Ki-67 | negative regulation of DNA endoreduplication | Proliferative marker present during all phases of the cell cycle except G0 | Tafra et al, 2014 |
| Smc1a | Structural maintenance of chromosomes 1a | negative regulation of DNA endoreduplication | Part of cohesin complex. Cohesin complex necessary for cohesion of sister chromatids after DNA replication and necessary for DNA double-strand break repair; may facilitate cell proliferation by preventing accumulation of DNA damage and ensuring genomic integrity during the process of cell cycle re-entry; Mutations in genes produce components of cohesins result in cohesinopathies i.e. Cornelia de Lange syndrome (CdLS). CdLS affects multiple organ systems (craniofacial, musculoskeletal, neurological, auditory, digestive tract) and intellectual function with sensorineural and conductive hearing loss being common | Gupta et al, 2016; Janek et al, 2016 |
| Smc3 | Structural maintenance of chromosomes 3 | negative regulation of DNA endoreduplication | See Smc1a description | See Smc1a description |
| Rad21 | Rad21 cohesin complex component | negative regulation of DNA endoreduplication | See Smc1a description | See Smc1a description |
| Stag1 | Stromal antigen 1 | negative regulation of DNA endoreduplication | See Smc1a description | See Smc1a description |
| Stag2 | Stromal antigen 2 | negative regulation of DNA endoreduplication | See Smc1a description | See Smc1a description |
| Shc1 | Shc adaptor  protein 1 | negative regulation of DNA endoreduplication | 3 isoforms (p66Shc, p52Shc, p46Shc); p66Shc participates in mitochondrial reactive oxygen species generation; critical for initiation of apoptosis by stress-activated p53; p52Shc and p46Shc may initiate Ras signaling | Wu et al, 2012 |

Selected References:

Froehlich, E.V., Rinner, B., Deutsch, A.J., Meditz, K., Knausz, H., Troppan, K., Scheipl, S., Wibmer, C., Leithner, A., Liegl, B., Lohberger, B. (2015) Examination of surviving expression in 50 chordoma specimens—A histological and in vitro study. J Orthop Res *33(5)*, 771-8.

Habtemichael, N., Heinrich, U.R., Knauer, S.K., Schmidtmann, I., Bier, C., Docter, D., Brochhausen, C., Helling, K., Brieger, J., Stauber, R.H., Mann, W.J. (2010) Expression analysis suggests a potential cytoprotective role of Birc5 in the inner ear. Mol Cell Neurosci *45(3)*, 297-305.

Chen, J., Wang, F., Gao, X., Zha, D., Xue, T., Cheng, X., Zhong, C., Han, Y., Qiu, J. (2009) Decreased level of cyclin A2 in rat cochlea development and cochlear stem cell differentiation. Neurosci Lett *453(3)*, 166-9.

Zhong, C., Han, Y., Ma, J., Zhang, X., Sun, M., Wang, Y., Chen, J., Mi, W., Xu, X., Qiu, J. (2015) Viral-mediated expression of c-Myc and cyclin A2 induces cochlear progenitor cell proliferation. Neurosci Lett *591*, 93-98.

Lu, N., Chen, Y., Wang, Z., Chen, G., Lin, Q., Chen, Z.Y., Li, H. (2013) Sonic hedgehog initiates cochlear hair cell regeneration through downregulation of retinoblastoma protein. Biochem Biophys Res Commun *430(2)*, 700-5.

Sherr, C.J. (1995) D-type cyclins. Trends Biochem Sci *20(5)*, 187-90.

Lee, S.H., Wang, X., Kim, S.H., Kim, Y., Rodriguez-Puebla, M.L. (2017) Cyclin D3 deficiency inhibits skin tumor development, but does not affect normal keratinocyte proliferation. Oncol Lett *14(3)*, 2723-2734.

Laine, H., Sulg, M., Kirjavainen, A., Pirvola, U. (2010) Cell cycle regulation in the inner ear sensory epithelia: role of cyclin D1 and cyclin-dependent kinase inhibitors. Dev Biol *337(1)*, 134-46.

Wang, H., Chen, H., Zhou, H., Yu, W., Lu, Z. (2017) Cyclin-dependent kinase inhibitor 3 promotes cancer cell proliferation and tumorigenesis in nasopharyngeal carcinoma by targeting p27. Oncol Res *25*, 1431-40.

Pavlides, S.C., Lecanda, J., Daubriac, J., Pandya, U.M., Gama, P., Blank, S., Mittal, K., Shukla, P., Gold, L.I. (2016) TGF-β activates APC through Cdh1 binding for Cks1 and Skp2 proteosomal destruction stabilizing p27kip1 for normal endometrial growth. Cell Cycle *15(7)*, 931-47.

Blaess, S., Graus-Porta, D., Belvindrah, R., Radakovits, R., Pons, S., Littlewood-Evans, A., Senften, M., Guo, H., Li, Y., Miner, J.H., Reichardt, L.F., Müller, U. (2004) Beta1-integrins are critical for cerebellar granule cell precursor proliferation. J Neurosci *24(13)*, 3402-12.

Kurima, K., Hertzano, R., Gavrilova, O., Monahan, K., Shpargel, K.B., Nadaraja, G., Kawashima, Y., Lee, K.Y., Ito, T., Higashi, Y., Eisenman, D.J., Strome, S.E., Griffith, A.J. (2011) A noncoding point mutation in Zeb1 causes multiple developmental malformations and obesity in Twirler mice. PLoS Genet *7(9)*, e1002307.

Tanno, B., Sesti, F., Cesi, V., Bossi, G., Ferrari-Amorotti, G., Bussolari, R., Tirindelli, D., Calabretta, B., Raschella, G. (2010) Expression of Slug is regulated by c-Myb and is required for invasion and bone marrow homing of cancer cells of different origin. J Biol Chem 285, 29434-29445

Ohashi, K., Togawa, T., Sugiura, T., Ito, K., Endo, T., Aoyama, K., Negishi, Y., Kudo, T., Ito, R., Saitoh, S. (2017) Combined genetic analyses can achieve efficient diagnostic yields for subjects with Alagille syndrome and incomplete Alagille syndrome. Acta Paediatr *106(11)*, 1817-1824.

Teng, C.S., Yen, H.Y., Barske, L., Smith, B., Llamas, J., Segil, N., Go, J., Sanchez-Lara, P.A., Maxson, R.E., Crump, J.G. (2017) Requirements for Jagged1-Notch2 signaling in patterning the bones of the mouse and human middle ear. Sci Rep *7(1)*, 2497.

Lewis, A.K., Frantz, G.D., Carpenter, D.A., de Sauvage, F.J., Gao, W.Q. (1998) Distinct expression patterns of notch family receptors and ligands during development of the mammalian inner ear. Mech Dev *78(1-2)*, 159-63.

Deng, L., Lu, Y., Zhao, X., Sun, Y., Shi, Y., Fan, H., Liu, C., Zhou, J., Nie, Y., Wu, K., Fan, D., Guo, X. (2013) Ran GTPase protein promotes human pancreatic cancer proliferation by deregulating the expression of surviving and cell cycle proteins. Biochem Biophys Res Commun *440(2)*, 322-9.

Yu, Y., Weber, T., Yamashita, T., Liu, Z., Valentine, M.B., Cox, B.C., Zuo, J. (2010) In vivo proliferation of postmitotic cochlear supporting cells by acute ablation of the retinoblastoma protein in neonatal mice. J Neurosci *30(17)*, 5927-36.

Rocha-Sanchez, S.M., Scheetz, L.R., Contreras, M., Weston, M.D., Korte, M., McGee, J., Walsh, E.J. (2011) Mature mice lacking Rbl2/p130 gene have supernumerary inner ear hair cells and supporting cells. J Neurosci *31(24)*, 8883-93.

Rubin, C.I., Atweh, G.F. (2004) The role of stathmin in the regulation of the cell cycle. J Cell Biochm *93(2)*, 242-50.

Jiang, W., Huang, S., Song, L., Wang, Z. (2018) STMN1, a prognostic predictor of esophageal squamous cell carincoma, is a marker of the activation of the PI3K pathway. Oncol Rep 39(2), 834-842.

Hitchens, M.R., Robbins, P.D. (2003) The role of the transcription factor DP in apoptosis. Apoptosis *8(5)*, 461-8.

Lin, Y.S., Chen, Y.J., Cohen, S.N., Cheng, T.H. (2013) Identification of TSG101 functional domains and p21 loci required for TSG101-mediated p21 gene regulation. PLoS One *8(11)*, e79674.

Chen, P., Zindy, F., Abdala, C., Liu, F., Li, X., Roussel, M.F., Segil, N. (2003) Progressive hearing loss in mice lacking the cyclin-dependent kinase inhibitor Ink4d. Nat Cell Biol *5(5)*, 422-6.

Malgrange, B., Knockaert, M., Belachew, S., Nguyen, L., Moonen, G., Meijer, L., Lefebvre, P.P. (2003) The inhibition of cyclin-dependent kinases induces differentiation of supernumerary hair cells and Deiters’ cells in the developing organ of Corti. FASEB J *17(14)*, 2136-8.

Laine, H., Sulg, M., Kirjavainen, A., Pirvola, U. (2010) Cell cycle regulation in the inner ear sensory epithelia: role of cyclin D1 and cyclin-dependent kinase inhibitors. Dev Biol *337(1)*, 134-46.

Oesterle, E.C., Chien, W.M., Campbell, S., Nellimarla, P., Fero, M.L. (2011) p27(Kip1) is required to maintain proliferative quiescence in the adult cochlea and pituitary. Cell Cycle *10(8)*, 1237-48.

Hu, B.H., Cai, Q., Manohar, S., Jiang, H., Ding, D., Coling, D.E., Zheng, G., Salvi, R. (2009) Differential expression of apoptosis-related genes in the cochlea of noise-exposed rats. Neuroscience *161(3)*, 915-25.

Li, L., Chen, K., Wu, Y., Long, Q., Zhao, D., Ma, B., Pei, D., Liu, X. (2017) Gadd45a opens up the promoter regions of miR-295 facilitating pluripotency induction. Cell Death Dis *8(10)*, e3107.

Marks, D.H., Thomas, R., Chin, Y., Shah, R., Khoo, C., Benezra, R. (2017) Mad2 overexpression uncovers a critical role for TRIP13 in mitotic exit. Cell Rep *19(9)*, 1832-1845. doi: 10.1016/j.celrep.2017.05.021.

Zhai, Y., Li, N., Jiang, H., Huang, X., Gao, N., Tye, B.K. (2017) Unique roles of the non-identical MCM subunits in DNA replication licensing. Mol Cell 67(2), 168-179. doi: 10.1016/j.molcel.2017.06.016.

Gao, J., Wang, Q., Dong, C., Chen, S., Qi, Y., Liu, Y. (2015) Whole exome sequencing identified MCM2 as a novel causative gene for autosomal dominant nonsyndromic deafness in a Chinese family. PLoS One 10(7), e0133522. doi: 10.1371/journal.pone.0133522. eCollection 2015.

Saadatzadeh, M.R., Elmi, A.N., Pandya, P.H., Bijangi-Vishehsaraei, K., Ding, J., Stamatkin, C.W., Cohen-Gadol, A.A., Pollok, K.E. (2017) The role of MDM2 in promoting genome stability versus instability. Int J Mol Sci 18(10), pii: E2216. doi: 10.3390/ijms18102216.

Laos, M., Sulg, M., Herranen, A., Anttonen, T., Pirvola, U. (2017) Indispensable role of Mdm2/p53 interaction during the embryonic and postnatal inner ear development. Sci Rep *7*, 42216. doi: 10.1038/srep42216.

Tafra, R., Brakus, S.M., Vukojevic, K., Kablar, B., Colovic, Z., Saraga-Babic, M. (2014) Interplay of proliferation and proapoptotic and antiapoptotic factors is revealed in the early human inner ear development. Otol Neurotol 35(4), 695-703.

Gupta, P., Lavagnolli, T., Mira-Bontenbal, H., Fisher, A.G., Merkenschlager, M. (2016) Cohesin’s role in pluripotency and reprogramming. Cell Cycle *15(3)*, 324-30.

Janek, K.C., Smith, D.F., Kline, A.D., Benke, J.R., Chen, M.L., Kimball, A., Ishman, S.L. (2016) Improvement in hearing loss over time in Cornelia de Lange syndrome. Int J Pediatr Otorhinolaryngol *87*, 203-7.

Wu, L., Sun, Y., Hu, Y.J., Yang, Y., Yao, L.L., Zhou, X.X., Wang, H., Zhang, R., Huang, X., Kong, W.J. (2012) Increased p66Shc in the inner ear of D-galactose-induced aging mice with accumulation of mitochondrial DNA 3873-bp deletion: p66Shc and mtDNA damage in the inner ear during aging. PLoS One *7(11)*, e50483.
